# Genetically targeted reporter imaging of deep neuronal network in the mammalian brain

**DOI:** 10.1101/2020.04.08.032870

**Authors:** Masafumi Shimojo, Maiko Ono, Hiroyuki Takuwa, Koki Mimura, Yuji Nagai, Masayuki Fujinaga, Tatsuya Kikuchi, Maki Okada, Chie Seki, Masaki Tokunaga, Jun Maeda, Yuhei Takado, Manami Takahashi, Takeharu Minamihisamatsu, Ming-Rong Zhang, Yutaka Tomita, Norihiro Suzuki, Anton Maximov, Tetsuya Suhara, Takafumi Minamimoto, Naruhiko Sahara, Makoto Higuchi

## Abstract

Positron Emission Tomography (PET) allows biomolecular tracking, while PET monitoring of brain networks has been hampered by the lack of a suitable reporter. Here, we describe *in vivo* brain imaging that takes advantage of bacterial dihydrofolate reductase, ecDHFR, and its unique antagonist, TMP. In mice, peripheral administration of radiofluorinated and fluorescent TMP analogs enabled PET and intravital microscopy, respectively, of neuronal ecDHFR expressions. This technique is applicable to the visualization of neuronal ensemble activities elicited by chemogenetic manipulation in the mouse hippocampus. Notably, ecDHFR-PET offers mapping of neuronal projections in non-human primate brains, indicating the availability of ecDHFR-based tracking technologies for network monitoring. Finally, we demonstrate the utility of TMP analogs for PET assays of turnover and self-assembly of proteins tagged with ecDHFR mutants. Our findings may facilitate a broad spectrum of PET analyses of a mammalian brain circuit at molecular levels that were not previously applicable for technical reasons.

## Introduction

Contemporary methods for live imaging of genetically-encoded reporters have transformed basic and translational neuroscience (*1, 2*). Many excellent fluorescence tools have recently been introduced for the visualization of neuronal ensembles and individual synapses, and for understanding the roles of neuronal cells in circuit development, function, and dysfunction (*3–5*). Despite its tremendous utility, fluorescence imaging remains largely unsuitable for simultaneous non-invasive monitoring of reporters in deep brain regions or across the entire brain because of light scattering and relatively narrow fields of view. One promising strategy for solving this technical bottleneck is to complement optical microscopy with positron emission tomography (PET). While PET itself has a limited spatiotemporal resolution, a combination of the two techniques may offer comprehensive insights into the dynamics of molecules of interest on scales ranging from subcellular to global levels (*6, 7*). However, no reports currently exist regarding bimodal imaging of the central nervous system with an intact blood-brain barrier (BBB).

A major challenge for such a bimodal analysis of exogenous markers genetically targeted in the brain is the development of selective radioactive ligands that efficiently penetrate the BBB (*8*). Indeed, even for the best-characterized PET reporter, herpes simplex virus 1 (HSV-TK), no BBB-permeable compounds have been identified to date (*9, 10*). Several groups have attempted PET with mutant forms of dopamine and cannabinoid receptors, but these methods have suffered from poor signal-to-noise ratio due to ligand interaction with endogenous proteins (*11, 12*). A recent study reported brain PET imaging with an isoform of pyruvate kinase, PKM2, and its associated radiotracer [^18^F]DASA-23 (*13*). Our laboratory has also demonstrated that a designer receptor exclusively activated by designer drugs (DREADD) and its artificial ligands, such as clozapine-N-oxide (CNO) and deschloroclozapine (DCZ), labeled with ^11^C could be employed for PET of neurons and transplanted iPS cells in animal brains (*14–16*). Nonetheless, these compounds cannot be applied to bimodal imaging, and they may also suffer from undesired effects on endogenous substrates (*13, 17*).

E. coli dihydrofolate reductase (ecDHFR) is an enzyme whose activity is specifically blocked by a small antibiotic, trimethoprim (TMP). TMP rapidly diffuses through tissues, passes the BBB and, importantly, lacks native targets in mammals (*18, 19*). Moreover, TMP can be conjugated with fluorophores, and its radioactive analogs, [^11^C]TMP and [^18^F]TMP, are compatible with PET (*20–23*). These features make ecDHFR and TMP attractive candidates for the design of versatile probes for PET and fluorescent imaging of intact brains. Here, we construct the first generation of such reporters and characterize their performance in brains of mice and non-human primates.

## Results

### *In vivo* visualization of ecDHFR reporter

We hypothesized that ecDHFR, along with TMP, could be a suitable candidate for bimodal optical and PET imaging of the neuronal circuits and their function in intact brains (**Figure 1A**). To explore this possibility, we first examined the utility of a red fluorophore-conjugated TMP derivative, TMP-hexachlorofluorescein (TMP-HEX), as a probe for optical imaging of ecDHFR in the brains of living mice. We locally introduced ecDHFR-EGFP into the mouse somatosensory cortex by an adeno-associated virus (AAV) vector under control of a pan-neuronal synapsin promoter (**Figure S1A, S1C**). Awake animals were examined by intravital two-photon laser microscopy after a single bolus tail vein administration of TMP-HEX. Individual neurons carrying ecDHFR-EGFP retained TMP-HEX, indicating that the compound penetrates the BBB similarly to conventional TMP (**Figure 1B**). Importantly, intraperitoneal administration of a saturating dose of unlabeled TMP prior to the imaging session almost completely blocked the binding of TMP-HEX to ecDHFR, supporting the specificity of TMP-HEX for ecDHFR *in vivo* (**Figure 1B, 1C**).

**Figure 1.**
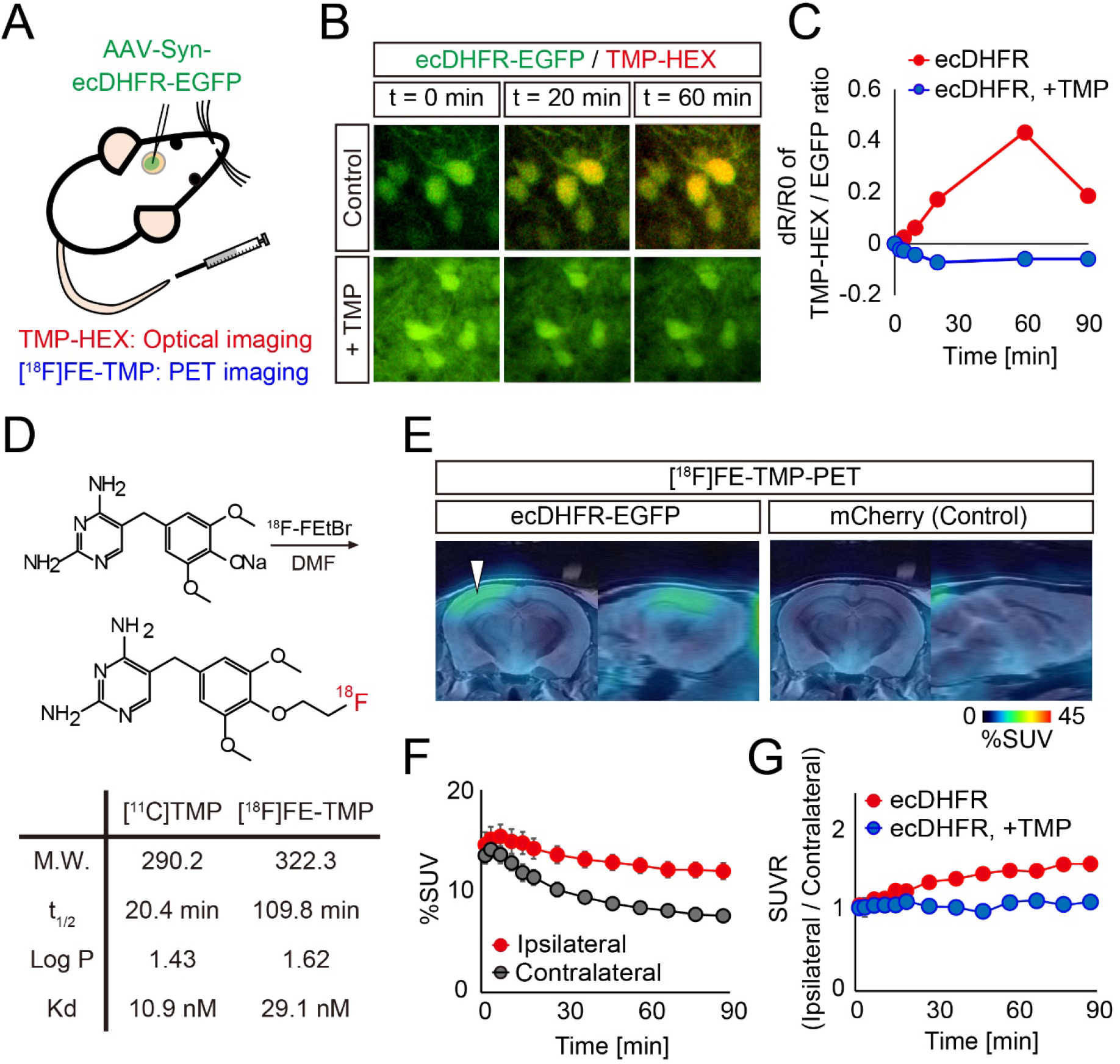
*In vivo* visualization of ecDHFR with TMP-HEX and [^18^F]FE-TMP. **A)** Schematic illustration of bimodal optical and PET reporter imaging in the brain of living mice. Recombinant ecDHFR-EGFP reporter proteins were introduced into the somatosensory cortex with AAV. Two-photon imaging and PET scan were conducted after intravenous administration of TMP analogs, TMP-conjugated fluorescence dye (TMP-HEX), and TMP labeled with radioisotope ([^18^F]FE-TMP), respectively. **B-C)** Peripherally delivered TMP-HEX penetrates the BBB and labels exogenously expressed ecDHFR in the mouse brain *in vivo*. Brains of awake animals were imaged under a two-photon microscope following tail vein administration of TMP-HEX (500 μg/kg). **B)** Typical merged images of EGFP (green) and TMP-HEX (red) fluorescence in mice that were given TMP-HEX alone (Control) or together with conventional TMP (i.p., 100 mg/kg). Each image is an average of 10 frames from serial z-stacks. **C)** Ratios of TMP-HEX fluorescence intensity relative to EGFP, plotted as a function of time. Averaged values from two independent experiments (20 cells/2 mice for each condition) are shown. **D)** Chemical structure and physical parameters of [^18^F]FE-TMP compared to conventional [^11^C]TMP. **E-G)** AAVs encoding ecDHFR-EGFP or mCherry (as control) were unilaterally injected into the somatosensory cortex. PET scans were performed after peripheral administration of radioactive TMP analog, [^18^F]FE-TMP alone, or [^18^F]FE-TMP together with conventional TMP (100 mg/kg). **E)** Representative PET images (coronal and sagittal sections) generated by averaging dynamic scan data at 60-90 min after i.v. injection of [^18^F]FE-TMP. Template MRI images were overlaid for spatial alignment. Arrowhead indicates the areas of accumulation of the radioactive ligand in animals carrying ecDHFR-EGFP (left). **F)** [^18^F]FE-TMP labeling kinetics in the brain of ecDHFR-EGFP-expressing mice (Mean ± SEM, n = 16). F(1, 30) = 8.753; p < 0.01 (two-way, repeated measure ANOVA). VOIs were manually placed on ipsilateral and contralateral areas for quantification. **G)** SUVR (%SUV ratio) of ipsilateral/contralateral signals in the brain of ecDHFR-EGFP-expressing mice after bolus injection of [^18^F]FE-TMP alone (n = 16) or together with conventional TMP (i.p., 100 mg/kg, n = 4). Data are plotted as Mean ± SEM. F(1, 18) = 6.896; p < 0.05 (two-way, repeated measure ANOVA).

To accomplish an *in vivo* macroscopic analysis of ecDHFR distribution in the brain by PET, [^18^F]fluoroethoxy-TMP ([^18^F]FE-TMP) (**Figure 1D**), a high-affinity ^18^F-labeled TMP derivative that has a prolonged half-life in radioactivity compared to conventional [^11^C]TMP (**Figure S2**), was next synthesized and injected into the tail vein of mice expressing ecDHFR-EGFP or red fluorescent protein, mCherry, as a control, followed by dynamic PET scans. Remarkably, mice expressing ecDHFR-EGFP exhibited strong radiosignals on an ipsilateral side of the somatosensory cortex locally delivered with AAV encoding ecDHFR-EGFP, but not in uninfected areas of the contralateral side or ipsilateral side expressing mCherry (**Figure 1E, 1F**). Pre-administration of unlabeled TMP significantly attenuated the enriched radioactive signals of [^18^F]FE-TMP on the ipsilateral side expressing ecDHFR-EGFP **(Figure 1G)**, indicating saturable and specific binding of [^18^F]FE-TMP to ecDHFR. Taken together, these results demonstrate that TMP analogs can be used for multiscale optical and PET imaging of ecDHFR-based reporters in the brains of live animals.

### PET monitoring of neuronal ensemble activities

To take advantage of the advanced utility of ecDHFR reporter for PET monitoring of neural circuit activations, we incorporated robust activity marking (RAM) system, an optimized synthetic activity-regulated element with the immediate early gene (IEG) cFos minimal promoter, which can drive the expression of a reporter protein in response to sensory stimuli and epileptic seizure (*24*) (**Figure 2A**). RAM-dependent efficient marking of neuronal activation was successfully validated in cultured neurons (**Figure S3A**). We then co-introduced AAVs encoding ecDHFR-d2Venus with RAM promoter and the excitatory DREADD, hM3Dq, into one side of the somatosensory cortex and examined the time-course change of the reporter expression by two-photon laser microscopy after air-puff stimulation of whiskers or hM3Dq activation via an intraperitoneal bolus administration of CNO. Repetitive sensory inputs by air-puff stimulation of whiskers robustly enhanced the fluorescence signals of ecDHFR-d2Venus in neurons, whereas the activation of hM3Dq more prominently induced signal enhancement in neurons expressing ecDHFR-d2Venus at 24 h after systemic administration of CNO (**Figure 2B, 2C**). We then prepared mice co-expressing RAM-ecDHFR-d2Venus and hM3Dq on one side of hippocampi, and tested whether PET can successfully capture CNO-provoked activations of a neural circuit. Remarkably, sequential PET scans with [^18^F]FE-TMP demonstrated significant accumulation of [^18^F]FE-TMP radiosignals at 24 h after injection of CNO (**Figure 2D, 2E**). A postmortem analysis of brain tissues collected from these mice following PET experiments demonstrated induced expression of ecDHFR-d2Venus primarily in the hippocampal dentate gyrus and CA3 neurons (**Figure S3B),** and there was good correlation between PET signals and reporter expressions (**Figure 2E**). These data indicate that [^18^F]FE-TMP PET successfully captured the chemogenetic activation of the hippocampal network with high sensitivity and specificity.

**Figure 2.**
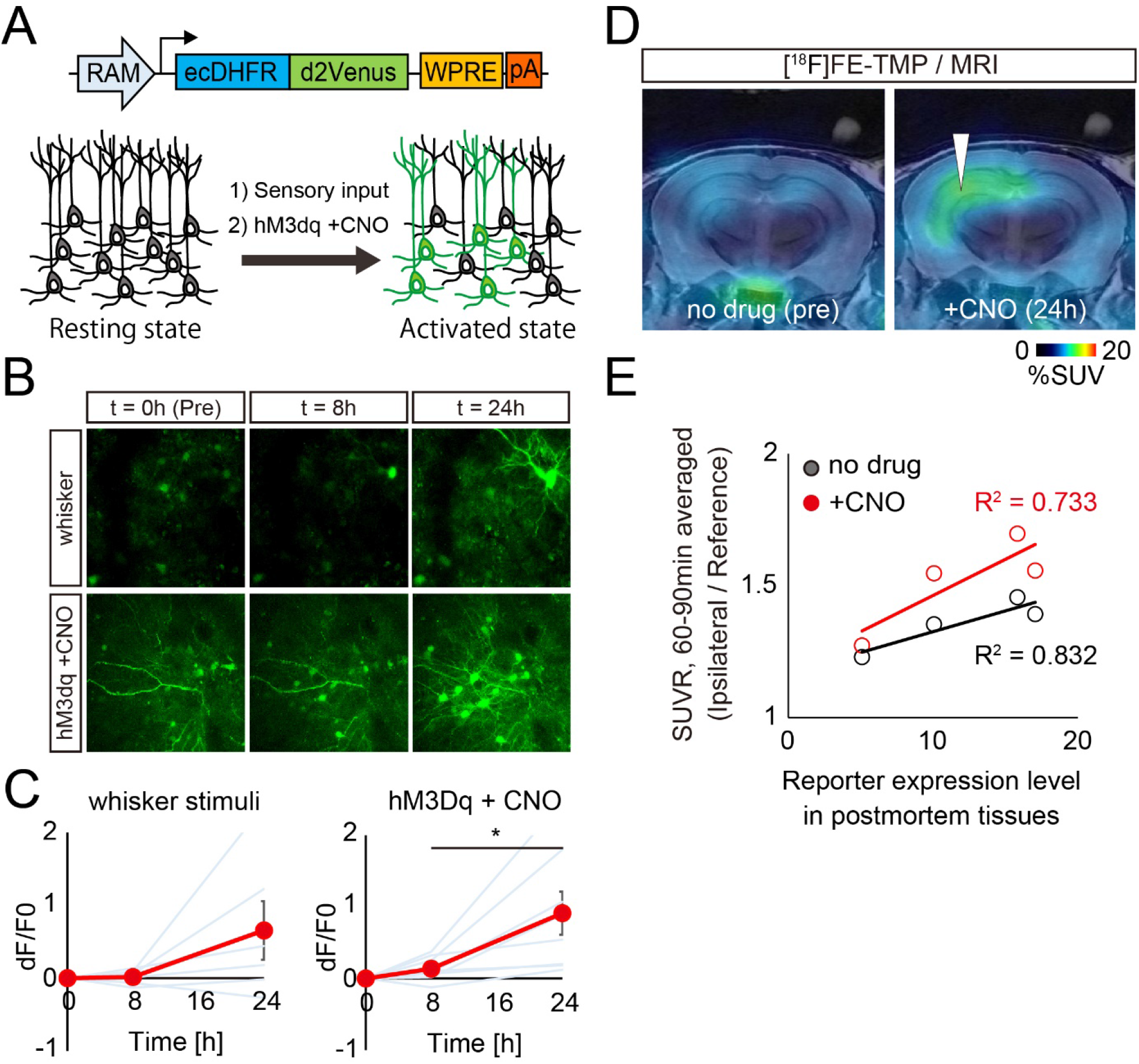
*In vivo* visualization of neuronal ensemble activation in response to sensory stimuli and hM3Dq-DREADD. **A)** Schematic illustration of virus construction and marking of neuronal ensemble activation by CNO mediated hM3Dq-DREADD activation. b-c, AAVs encoding ecDHFR fused to destabilized Venus (ecDHFR-d2Venus) with RAM promoter and hM3Dq were co-introduced into two segregated regions of the somatosensory cortex and imaged under two-photon microscope. **B)** Fluorescent intensity of ecDHFR-d2Venus in the brain of awake animals was analyzed before (0 min) and after (8 h or 24 h) air-puff mediated whisker stimulation (10 sec, 5 times) or chemogenetic activation of hM3Dq via peripheral administration of CNO (10 mg/kg). Averaged images of 10 stacked frames in serial z position are shown. **C)** Averaged ecDHFR-d2Venus fluorescence intensities, plotted as dF/F0 ratios at different time points after neuronal activation. Data from 6-8 regions / 3-4 mice for each condition are plotted as Mean ± SD (red line) and individual regions (blue line). 8 h vs 24 h, F(2, 21) = 7.715; *p < 0.05 (one-way ANOVA). **D-E)** Mice expressing ecDHFR-d2Venus and hM3Dq in a side of hippocampus was sequentially analyzed by [^18^F]FE-TMP PET imaging before (left) or 24 h after (right) intraperitoneal CNO injection (0.3 mg/kg). **D)** Representative PET images demonstrate that 0.3 mg/kg CNO-mediated activation of hM3Dq enhanced the accumulation of radioactive signals in the hippocampus. Averaged images of dynamic scan data at 60-90 min after i.v. injection of [^18^F]FE-TMP are shown. Template MRI images were overlaid for spatial alignment. Arrowhead indicates selective accumulation of radioactive signals in the AAV injection site after CNO administration. **E)** SUVR (%SUV ratio) of ipsilateral hippocampi / reference region (brain stem) signals during dynamic PET scans. Data from ecDHFR-d2Venus expressing mice (n = 4) before and after 0.3 mg/kg CNO i.p. injection are plotted. Correlation with R^2^ values between SUVR of [^18^F]FE-TMP_PET and expression levels of ecDHFR-d2Venus (fluorescence) in postmortem tissues was determined by linear regression analysis.

### *In vivo* illumination of deep neural circuit

We next sought to determine whether our imaging techniques can be leveraged to trace neural circuits in deep brain regions of live non-human primates in more detail than rodents. Given the similarity of anatomies of monkey and human brains, we reasoned that this approach could further validate the current method for future medical applications. An AAV encoding ecDHFR-EGFP was delivered to one side of the neocortex and striatum of a 3.4-year-old male common marmoset to express the reporter in the telencephalon. PET scans of this animal with [^18^F]FE-TMP were performed 45 days after virus injection. To our surprise, the PET scans revealed robust accumulation of [^18^F]FE-TMP not only in brain regions proximal to the injection site (i. e., neocortex, caudate nucleus, and putamen) but also in other spatially segregated but interconnected areas, including the thalamus and midbrain substantia nigra (**Figure 3A**). The marmoset brain displayed more efficient radioligand uptake than the mouse brain, and the putamen-to-cerebellum ratio of radioactive signals increased up to 4 at the end of the dynamic PET scan (**Figure 3B**). Again, distribution of the radioactive signal was consistent with the localization of transgene expression, as assessed by postmortem microscopic imaging of brain sections (**Figure 3C**). It is accordingly likely that the current ecDHFR-based reporters, particularly in combination with the RAM system, are capable of pursuing chemogenetic activation of a specific neural connection in non-human primates, which has been technically enabled by DREADD and DCZ in our recent work (*16*).

**Figure 3.**
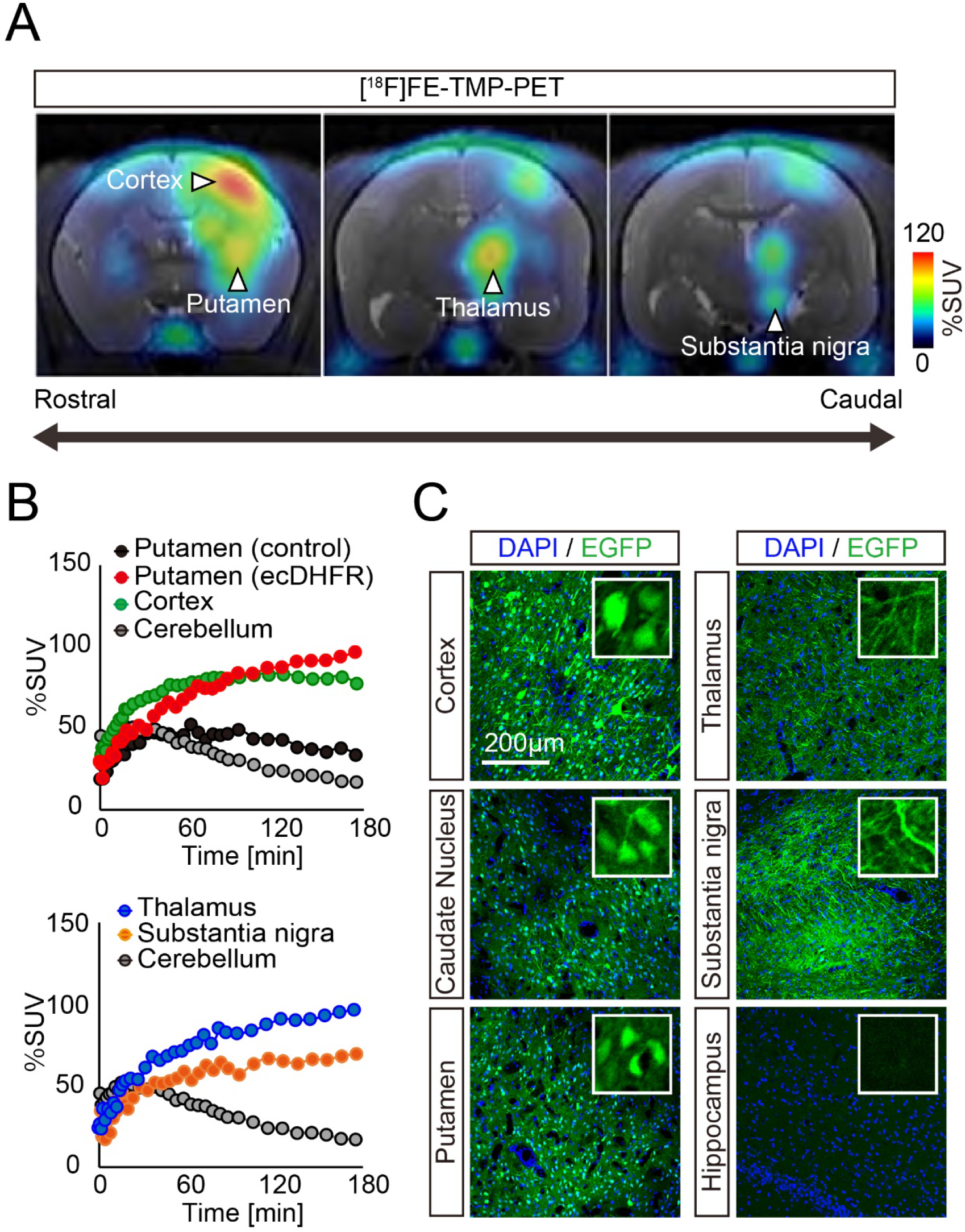
PET imaging of ecDHFR/TMP reporters in primate brain. ecDHFR-EGFP was expressed in the brain of a 3.4-year-old common marmoset with an AAV. PET scans of [^18^F]FE-TMP were performed 45 days after virus injection. **A)** Representative coronal PET images generated by averaging dynamic scan data at 60 - 180 min after i.v. injection of [^18^F]FE-TMP (left). Note that reporter molecule was densely distributed in thalamus and substantia nigra pars compacta, which are connected to the neocortex and putamen via direct neuronal tract, respectively. PET images are overlaid with an MRI template. Scale bar represents %SUV. **B)** Time-radioactivity curves for [^18^F]FE-TMP in the putamen carrying ecDHFR-EGFP (red symbols) or control AAV (black symbols), neocortex (green symbols), and cerebellum (filled gray symbols) are displayed in the upper panel. Curves in the thalamus (blue symbols) and substantia nigra (orange symbols) along with cerebellar data are also shown in the lower panel. **C)** Postmortem analysis of ecDHFR-EGFP expression in brain slices. Representative images illustrate EGFP fluorescence in the cortex, caudate nucleus, putamen, thalamus, and substantia nigra. High magnification image frames are shown in inserts.

### PET analysis of protein turnover with ecDHFR mutant

In the final set of experiments, we aimed to explore further applications of unique ecDHFR mutants for PET assays of the protein dynamics in the brain circuit. To this end, we employed an ecDHFR mutant with a destabilized domain (DD) which enabled us to control gene expression and function of a protein of interest (POI) fused to DD in a manner inducible by TMP administration (**Figure 4A, S4**) (*25, 26*). In agreement with previous studies, treatments with TMP rapidly restored the levels of DD-EGFP in the mouse brain (**Figure 4B, 4C**). We subsequently utilized a cyclic phosphodiesterase, PDE10A, as a POI, which can be detected by PET with a specific radioligand, [^18^F]MNI659 (*27*). The catalytic domain of human PDE10A (hPDE10A_CD, **Figure S5**) was incorporated in AAVs encoding hPDE10A_CD-EGFP with N-terminal ecDHFR or DD tags and was inoculated into the neocortex. PET scans demonstrated a significant accumulation of [^18^F]MNI659 in the ipsilateral area expressing DD-hPDE10A_CD-EGFP following peripheral delivery of TMP, suggesting that PET can be used to monitor acute drug-inducible stabilization of DD-POIs in a brain circuit (**Figure 4D, 4E**). Hence, the current technology would serve for manipulation of molecular signaling by switching POI on and off and for assessing the functionality of protein degradation systems in healthy and diseased neural networks.

**Figure 4.**
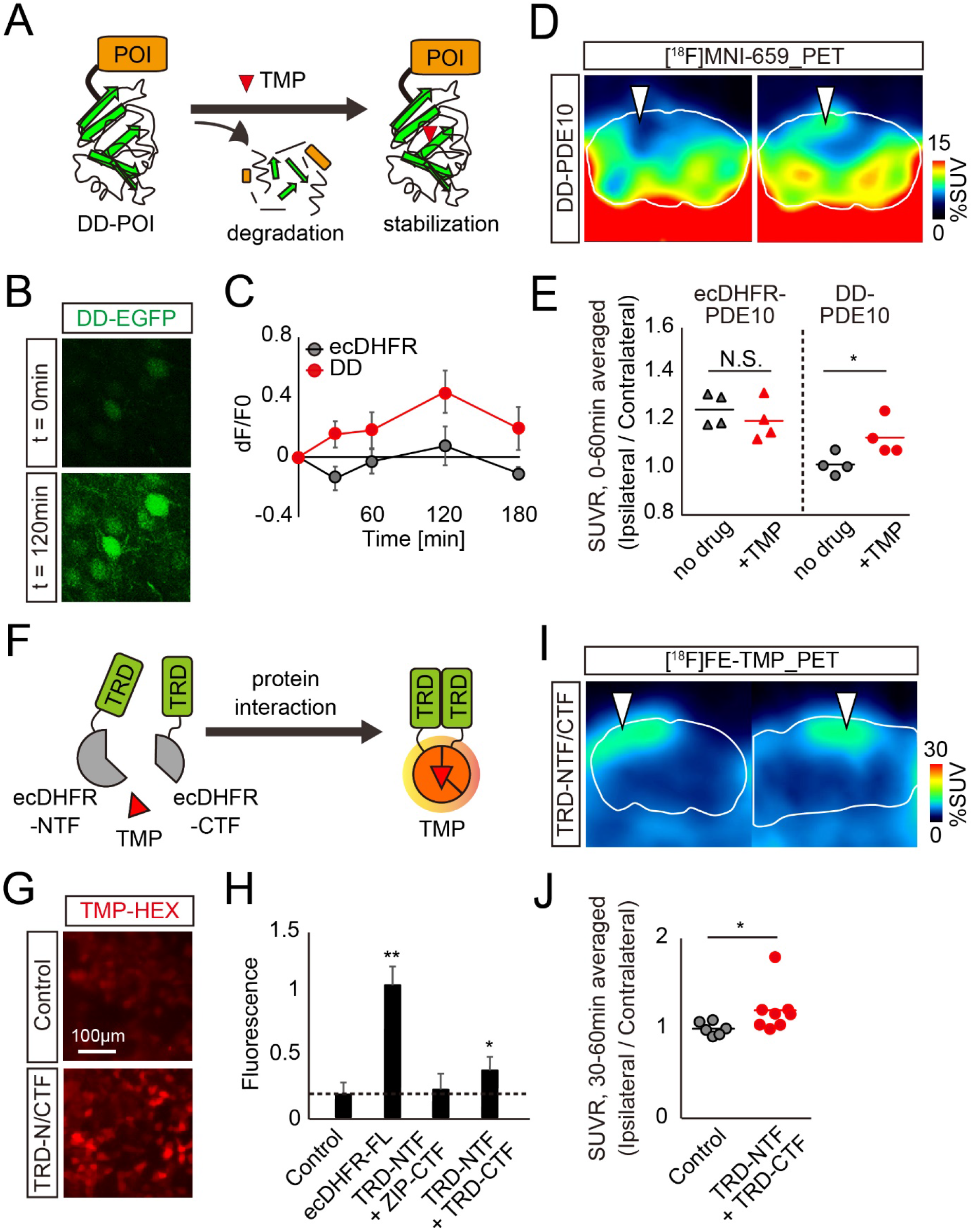
PET analysis of TMP-inducible protein stabilization and protein-protein interaction in the brain. **A)** Schematic illustration of TMP-dependent control of DD-POI protein turnover. A protein of interest (POI) fused to a destabilizing domain (DD) undergoes rapid degradation by ubiquitin-proteasome system, while POI can be stabilized in the presence of TMP. **B-C)** Mice expressing fusion proteins comprised of EGFP and either wild-type or destabilized mutant ecDHFR (ecDHFR-EGFP or DD-EGFP, respectively) were imaged by two-photon microscope. The constructs were introduced into the somatosensory cortex with AAVs. **B)** Fluorescent intensity of DD-EGFP in intact brains of awake animals was analyzed before (0 min) and after (120 min) peripheral administration of TMP (100 mg/kg). Averaged images of 10 stacked frames in serial z position are shown. **C)** Averaged ecDHFR-EGFP and DD-EGFP fluorescence intensities plotted as dF/F0 ratios at different time points after TMP delivery. Data from 10 neurons for each genotype are plotted as Mean ± SD. **D-E)** ecDHFR-PDE10 and DD-PDE10 were expressed in a caudal region of cortical hemisphere from an AAV. Accumulation of the radioactive signal was monitored by PET scan with [^18^F]MNI659, a selective radioactive PET ligand against PDE10. **D)** Representative PET images demonstrate the detection of TMP-stabilized DD-PDE10 in caudal area of the neocortex. Averaged images of dynamic scan data at 0-30 min after i.v. injection of [^18^F]MNI659 are shown. 90-min scans were conducted before or 2 h after intraperitoneal TMP injection (100 mg/kg). White lines mark the whole brain area as determined by MRI. Note selective accumulation of radioactive signals in the AAV injection site by TMP administration (arrowheads). **E)** Averaged SUVR (%SUV ratio) of ipsilateral/contralateral radioactive signals during 0-60 min scan in individual animals expressing ecDHFR-PDE10 (n = 4) or DD-PDE10 (n = 4). Data represent Mean (horizontal bar) and values from individual animals (dots) for each group under control condition (no drug) or after TMP treatment. t(6) = 2.521; *p < 0.05 (Student’s t-test). Note a significant TMP-dependent increase of the ratio in animals carrying a destabilized reporter. **F)** Schematic representation of a reporter containing the N- and C-terminal ecDHFR fragments (NTF and CTF). These fragments reassemble after interaction of fused proteins A and B, thereby restoring the TMP-binding ability. **G)** Representative images illustrate a selective retention of TMP-HEX in cells expressing ecDHFR NTF and CTF fused with human tau repeat domain (TRD). **H)** TRD oligomerization-dependent TMP-HEX labeling was quantitatively analyzed in cells expressing various combinations of ecDHFR-NTF and CTF fragments as normalized fluorescence intensities. Mean value of full-length ecDHFR (ecDHFR-FL) was set as 1. Data from seven independent experiments are shown as Mean ± SD. F(3, 24) = 77.86; **p < 0.01, *p < 0.05 (one-way ANOVA followed by Dunnett post-hoc test). **I-J)** PET analysis of Tau oligomerization in the brain. 90-min [^18^F]FE-TMP PET scans were conducted after co-injection of AAVs encoding TRD-NTF and TRD-CTF into one hemisphere of the somatosensory cortex. **I)** Representative PET images (coronal and sagittal sections from left) were generated by averaging dynamic scan data at 60-90 min after i.v. injection of [^18^F]FE-TMP. Images show a selective accumulation of radioactive signals in the AAV injection site (arrowheads). White lines mark the whole brain area as determined by template MR image. **J)** Averaged SUVR (%SUV ratio) of ipsilateral/contralateral radioactive signals during 15-60 min dynamic PET scans. Data from Control mice (n = 6; mixture of uninfected mice, n = 4 and TRD-NTF/ZIP-CTF-expressing mice, n = 2) or TRD-NTF/TRD-CTF-expressing mice (n = 8) are plotted as Mean (horizontal bar) and individual animals (dots). *p < 0.05 (Mann-Whitney test).

### PET analysis of protein complex formation

We also relied on a protein-fragment complementation assay (PCA) to clarify the usability of ecDHFR split mutants for monitoring the oligomerization of misfolded microtubule-associated protein tau, which may propagate through the neural network, leading to neurodegeneration in the pathological cascade of Alzheimer’s disease (*28, 29*). We constructed an ecDHFR-PCA system in which the N- and C-terminal fragments of the ecDHFR protein (NTF and CTF, respectively) reassemble, thereby restoring its capability to bind to TMP, through interaction between a bait and prey (**Figure 4F, S6A-S6D**). As a proof-of-principle experiment, TMP-HEX efficiently labeled cultured cells co-expressing NTF and CTF conjugated to a human tau repeat domain (TRD), suggesting that the self-assembly of TRD was spontaneously initiated and could be detected by our method *in vitro* (**Figure 4G, 4H**). We therefore co-injected AAVs encoding TRD-NTF and TRD-CTF in the somatosensory cortex, and conducted PET scans with [^18^F]FE-TMP to detect potential tau oligomerization and propagation *in vivo* more than one month after the surgical procedure. Remarkably, we found an accumulation of [^18^F]FE-TMP in the proximity of the inoculation site (**Figure 4I**), and an ipsilateral-to-contralateral ratio of PET signals in a steady state was significantly increased in brains of these mice relative to control brains (**Figure 4J**). A PET scan with [^11^C]PBB3, a potent PET tracer for fibrillary deposits of pathological tau proteins (*30*), did not reveal any significant accumulation of radioactive signals in the areas receiving the AAV delivery (**Figure S6E, S6F**). Hence, PET scans with [^18^F]FE-TMP could capture low-order, PBB3-undetectable tau assemblies, which potentially propagate along a specific brain circuit in the brains of intact animals.

## Discussion

In summary, we have designed and validated new ecDHFR-based reporters and their ligands for complementary fluorescence and PET analyses of the spatial distribution, stability, and aggregation of genetically-targeted fusion proteins in brain circuits of live animals. Importantly, neither overexpression of the reporter nor repetitive treatment with TMP affects neuronal physiology and animal behaviors (*25, 31*). Accordingly, our approaches enable high-contrast imaging and operations of a specific molecule at a network level in physiological and pathophysiological conditions, which is a task that could not be easily accomplished by conventional techniques. For instance, when driven by cell type-specific or activity-regulated promoters, ecDHFR reporters become instrumental for understanding how various biologically active molecules as represented by optogenetic and chemogenetic tools acutely impact the neural circuit structure and brain function in mice and non-human primates. In genetically engineered animals, our technology allows a longitudinal whole-brain assessment of cell type distribution, brain-wide circuit reorganization, and pharmacological effects on the circuit integrity in living animals. Moreover, ecDHFR-PCA could help to elucidate the mechanisms of neurodegenerative proteinopathies, as Holmes et al. have recently established a fluorescence resonance energy transfer-based assay to monitor the seeding ability of tau aggregates *in vitro* (*32*). In conjunction with currently available imaging modalities such as functional magnetic resonance imaging and functional near-infrared spectroscopy (*33*), our reporter methodologies will facilitate the construction of new assaying systems elucidating links between molecules and functions in a target network. Since the ecDHFR-TMP system is suitable for mammals, recent advances in the design of viral tools for non-invasive gene delivery make it feasible to eventually apply these reporters to biomedical PET imaging of human brains (*34*).

## Material and Methods

### Reagents

The following chemicals and antibodies were commercially purchased and used in this study: TMP (Sigma, T7883); TMP-Hexachlorofluorescein, TMP-HEX (Active Motif, 34104); chicken anti-GFP (AVES, GFP-1020); mouse anti-βActin (Sigma, A1978); mouse anti-FLAG, clone M2 (Sigma, F1804); mouse anti-HA (Covance, MMS-101R); rabbit anti-NeuN (Cell Signaling Tech, #12943S); rabbit anti-HA (Cell Signaling Tech, #3724); rabbit anti-cFos (Cell Signaling Tech, #2250); mouse anti-GFAP (Millipore, MAB360); rabbit anti-PDE10 (Protein Tech, 18078-1-AP); chicken anti-MAP2 (Abcam, AB5392).

### Animals

All procedures involving animals and their care were approved by the Animal Ethics Committee of the National Institutes for Quantum and Radiological Science and Technology. We used postnatal day 0 or 2 month-old C57BL/6j mice (Japan SLC) for the genetic manipulation by Adeno Associate Virus (AAV) mediated gene delivery followed by two-photon microscopy, PET scan, histology, and metabolite analysis. For *in vivo* injection of AAV into neonatal pups, animals were deeply anesthetized on ice, and 1.0 μL of purified AAV stocks (titer ranging from 0.5 x10^9^ to 2.0 x10^9^ viral genomes) was injected directly into the side of a lateral cerebral ventricle via glass capillary. This technique was established as a relatively simple and fast method to manipulate gene expression in extensive brain regions with minimal long-term damages (*35, 36*), and we indeed observed that mice receiving AAV injection grew normally and exhibited expression of constructs in regions adjacent to ventricles, including the retrosplenial cortex and hippocampus without obvious neuroanatomical abnormalities at 2 months of age (**Figure S1B**). We performed PET scans of these animals at 2 months of age. For AAV injection into adult mouse cortex, mice at 2-3 months of age were deeply anesthetized with 1.5-2.0% isoflurane and underwent a surgical procedure to attach cranial windows to one side of the somatosensory cortex (center was positioned 1.8 mm caudal and 2.5 mm lateral from bregma). 1.0-1.5 μL of AAV (titer ranging from 2.0 x10^9^ to 8.0 x10^9^ viral genomes) was slowly injected via glass capillary into parenchyma positioned at a depth of 0.250-0.375 mm from the cortical surface during this procedure, and this model allowed us to perform side-by-side comparative imaging within a single animal (**Figure S1A**). Alternatively, stereotactic injection was coordinated for AAV delivery into somatosensory cortex (stereotactic coordinates: anteroposterior, −1.5 mm; mediolateral, 2.0 mm; dorsoventral 0.3 mm; relative to bregma) or hippocampus (stereotactic coordinates: anteroposterior, −2.5 mm; mediolateral, 2.0 mm; dorsoventral 1.5 mm; relative to bregma). We performed two-photon microscopic and PET imaging more than one month after the surgical procedure. At this time point, intact BBB integrity was validated as radioactive signals of PET probes [^11^C]GF120918 and [^11^C]verapamil, well characterized substrates for P-glycoprotein (*37*), and did not accumulate at the AAV injection site and neighboring areas (**Figure S7**). All mice were housed with littermates (2-5 mice depending on litter size) and maintained in a 12 h light/dark cycle with ad libitum access to standard diet and water.

A 3.4-year-old male common marmoset was used as model for tracing the neuronal circuit in primate brain. The animal was born at the National Institutes for Quantum and Radiological Science and Technology and was housed as one of a pair, with one family member of the same sex, in a cage (0.66 (h)×0.33 (w)×0.6 (d) m). Environmental lighting was provided from 8:00 a.m. to 8:00 p.m. in this colony room. Room temperature was maintained at 24–30°C and relative humidity at 40–60%. Balanced marmoset food pellets (CMS-1, CLEA Japan, Tokyo, Japan) were provided at sufficient amounts once a day, and water was available ad libitum in their home cage. For AAV injection, the marmoset was immobilized by intramuscular (i.m.) injection of ketamine (25 mg/kg) and xylazine (0.2 mg/kg) and intubated with an endotracheal tube. Anesthesia was maintained with isoflurane (1-3%, to effect). The marmoset then underwent a surgical procedure to open burr holes (~5 mm diameter) for the injection needle. During surgery, CT images were captured and overlaid on magnetic resonance (MR) images using PMOD software (PMOD Technologies Ltd, Zurich Switzerland) to obtain stereotaxic coordinates for spatial alignment between the injection needle and target location. AAV encoding ecDHFR-EGFP expression cassette was slowly delivered (3 μL at 0.25 μL/min) into one side of the neocortex and striatum, and control AAV was injected into the other, by means of a microsyringe (NanoFil, WPI, Sarasota, FL, USA) with a 33-G beveled needle that was mounted into a motorized microinjector (Legato130, KD Scientific or UMP3T-2, WPI, Sarasota, FL, USA). Other surgical procedures and viral injection was as described previously (*15*).

### Radiosynthesis

[^11^C]TMP (trimethoprim) was radiosynthesized using the precursor compound 4-((2,4-diaminopyrimidin-5-yl)methyl)-2,6-dimethoxyphenol (Sundia). [^11^C]Methyl iodide was produced and transferred into 300 μl of N,N-dimethylformamide (DMF) containing 1 mg of precursor and 7.2 μl of 0.5 N aqueous sodium hydroxide at room temperature. The reaction mixture was heated to 80°C and maintained for 5 min. [^18^F]fluoroethoxy-TMP ([^18^F]FE-TMP) was radiosynthesized using the precursor compound sodium 4-((2,4-diaminopyrimidin-5-yl)methyl)-2,6-dimethoxyphenolate (Sundia). [^18^F]Fluoroethyl bromide was produced and transferred into 250 μl of DMF containing 1.5 mg of precursor at room temperature. The reaction mixture was heated to 100°C and maintained for 10 min. After cooling the reaction vessel, the radioactive mixture was transferred into a reservoir for HPLC purification (Atlantis Prep T3, 10 x 150 mm; [^11^C]TMP: methanol/50 mM ammonium acetate = 40/60, 5.0 ml/min; [^18^F]FE-TMP: methanol/50 mM ammonium acetate = 35/65, 4.5 ml/min; UV: 254 nm). The fraction corresponding to [^11^C]TMP or [^18^F]FE-TMP was collected in a flask containing 100 μl of 25% ascorbic acid solution and Tween 80 and was evaporated to dryness under a vacuum. The residue was dissolved in 3 ml of saline (pH 7.4) to obtain [^11^C]TMP or [^18^F]FE-TMP as an injectable solution. The final formulated products were radiochemically pure (95%) as determined by HPLC (Atlantis T3, 4.6 x 150 mm; methanol/50 mM ammonium acetate = 40/60, 1 ml/min; UV: 254 nm). Radiosynthesis of [^11^C]PBB3, a specific PET tracer for aggregated filamentous tau protein, was described previously (*30*). [^18^F]MNI659, a highly selective antagonist against phosphodiesterase 10A (PDE10A), was radiosynthesized by previously described methods (*38*). [^11^C]GF120918 and [^11^C]verapamil were radiosynthesized as described previously (*37*). Ethyl [^18^F]fluoroacetate was prepared in a manner similar to that reported (*39*). Briefly, an anhydrous acetonitrile solution of ethyl 2-(*p*-toluenesulfonyloxy)acetate (3 mg/500 μL, Tokyo Chemical Industry Co., Ltd, Tokyo, Japan) was added to a reaction vessel containing dry [K/Kryptofix 2.2.2]^+18^F^-^, and the reaction mixture was left standing for 10 min at 105°C. The reaction solution was then applied to HPLC (column, XBridge C18 (Waters Corporation; Milford, MA, USA), 5 μm, 10 i.d × 250 mm; mobile phase; saline/ethanol=90/10; flow rate, 4 mL/min (tR: ca. 8.0 min)). The fraction containing ethyl [^18^F]fluoroacetate was collected in an injection vial. Total reaction time after bombardment was around 45 min. The yield was 47 ± 7.2% (n = 3) at the end of synthesis, with a radiochemical purity of more than 99%.

### Plasmid construction and virus preparation

pFUGW lentivirus shuttle plasmids with human Ubiquitin C promoter with customized multi-cloning site were described previously (*40*). cDNAs of wild-type ecDHFR (ecDHFR) and leucine zipper motif of Saccharomyces cerevisiae GCN4 (235-281 aa) were synthesized as a gBlocks gene fragment (Integrated DNA Technologies). pBMN-ecDHFR(DD)-YFP was a kind gift from Dr. Thomas Wandless (Addgene plasmid #29325). pAAV-RAM-d2Venus plasmid was a kind gift from Drs. Tetsuo Yamomori and Masanari Ohtsuka (RIKEN). cDNA of Human Phosphodiesterase type 10 (hPDE10) was amplified by conventional nested PCR using human brain cDNA (Clontech). cDNAs template encoding wild-type and D674A mutant form of hPDE10 catalytic domain (CD, 449-789 aa), FKBP12-rapamycin binding domain (FRB), and FK506 binding protein (FKBP) were synthesized as gBlock gene fragments. cDNAs encoding full-length or repeat domain (RD) of human 2N4R tau protein were as described previously (*41*). For protein complementation assay of ecDHFR reporter, N-terminal fragment (NTF, 1-87 aa) or C-terminal fragment (CTF, 88-159 aa) of ecDHFR was conjugated to ZIP or human Tau repeat domain (TRD) via linker sequence of GGGGSGGGGS. To generate plasmid vectors for expression of fluorescent or fusion proteins including EGFP, tdTomato, d2Venus, ecDHFR-EGFP, ecDHFR-d2Venus, DD-EGFP, full-length hPDE10 746 aa variant, ecDHFR-hPDE10_CD-EGFP, ecDHFR-hPDE10(D674A)_CD-EGFP, DD-hPDE10_CD-EGFP, FLAG-ecDHFR-HA, ZIP-FLAG-ecDHFR(NTF), ZIP-ecDHFR(CTF)-HA, TRD-FLAG-ecDHFR(NTF), and TRD-ecDHFR(CTF)-HA, each cDNA fragment was amplified by PCR and reassembled in the multi-cloning site of pFUGW plasmid. Sequences of these plasmid DNAs were verified and directly used for transfection.

For production of recombinant lentiviruses, pFUGW shuttle vector, pVSVg, and pCMVΔ8.9 plasmids were co-introduced into human embryonic kidney 293T cells using the FuGENE reagent (Promega). 48 h after transfection, secreted viruses in medium were collected, filtered through a 0.45 μm filter, and infected to neuronal culture as described previously (*40*). Recombinant AAV plasmid with rat synapsin promoter (pAAV-Syn) followed by customized multi-cloning site, woodchuck posttranscriptional regulatory element (WPRE), and polyA signal flanked with ITRs was described previously (*26*). cDNAs described above were directly subcloned into the multi-cloning site of pAAV-Syn plasmid from pFUGW-based lentivirus plasmids. In vitro validation well confirmed the specific expression of EGFP in neurons from a locally injected AAV under the control of the pan-neuronal synapsin promoter (**Figure S1C**). For activity dependent marking of neuronal ensemble, Syn promoter was substituted with RAM promoter. AAV encoding hM3-Dq was described previously (*16*). All other AAVs were prepared in large scale HEK293T cell cultures that were transfected with mixture plasmids of AAV encoding full-length cDNAs and the serotype DJ packaging plasmids pHelper and pRC-DJ with polyethyleneimine (Polysciences). 48 h after transfection, cells were harvested, extracted, and then AAV particles were purified using HiTrap heparin column (GE Healthcare) as described previously (*42*). The virus titer of purified AAV stocks was determined by AAVpro^®^ Titration kit (for Real Time PCR) ver2 (TaKaRa).

### Cell Culture

HEK293T cells were grown and maintained in DMEM (Invitrogen) supplemented with 10% fetal bovine serum (Sigma) and Penicillin/Streptomycin (Invitrogen) at 37°C in a 5% CO_2_ incubator. For all transfection experiments, cells were plated in 24-well plates and transfected with 0.25 μg plasmid DNA and 1.25 μL FuGENE (Promega). Primary neuronal culture was prepared as described previously (*40*). Briefly, cortex and hippocampus from E17.5 mouse embryos were dissociated and plated onto poly-D-lysine (Sigma) coated 18 mm glass coverslip at a density of 200,000-250,000 cells/cm^2^. Neurons were grown for 3-4 days in Neurobasal medium (Thermo Fisher) supplemented with 2% FBS (HyClone), Glutamax (Thermo Fisher), and B27 (Thermo Fisher) and infected with recombinant lentivirus encoding d2Venus with RAM promoter, and further maintained in serum-free medium. At DIV21, medium was replaced with solution containing 10 mM HEPES-NaOH, pH7.4, 140 mM NaCl, 5 mM KCl, 0.8 mM MgCl_2_, 2 mM CaCl_2_ and 10 mM Glucose, and neurons were stimulated by addition of 15 mM KCl or 100 μM Picrotoxin (Tocris) for 6 h followed by fixation for immunocytochemical analysis as described previously (*43*). Fluorescence images were then acquired by FV1000 laser scanning confocal microscopy (Olympus) with x60 (NA 1.35) oil immersion objectives.

### Live cell imaging and immunoblot

To assess the stabilization of DD-EGFP with TMP analogs, HEK293T cell culture medium was replaced with fresh medium with or without 10 μM TMP (Sigma) or FE-TMP at 4 h after transfection, and further incubated for additional 24 h. The next day, images of EGFP signal were captured, and cells were lysed in 1x SDS-PAGE sample buffer in Tris-HCl, pH7.4, 150 mM NaCl, 2 mM EDTA for immunoblot analysis. The protein extracts were separated by either 10% or 10-20% Tris-glycine gel, transferred to a nitrocellulose membrane, and subsequently probed with primary antibodies and HRP-conjugated secondary antibodies (Jackson IR Lab). Chemiluminescent signal was elicited using ECL (PIERCE) reagents and detected by c-Digit blot scanner (LI-COR) or Amersham Imager 600 (GE Healthcare). Signal intensity was quantified by Image J software. For fluorescence time-lapse imaging, transfected HEK293T cells on 24-well plates were detached by trypsin and re-plated onto poly-D-lysine (Sigma) -coated 25-mm glass coverslips. 24 h after plating, the coverslips were transferred to a custom-built perfusion chamber mounted on the stage of an inverted Olympus IX83-based fluorescence microscope controlled by Olympus Cell Sense Dimension software. Time series images were acquired at 0.2 Hz for 15 min at room temperature with an Orca-Flash4.0 cMOS camera (Hamamatsu Photonics) using a x20 objective lens (NA 0.45). At the beginning and ending time points, EGFP images were captured as references of transgene expression in each cell. Cells were perfused with HEPES buffered solution (10 mM HEPES-NaOH, pH7.4, 140 mM NaCl, 5 mM KCl, 0.8 mM MgCl_2_, 2 mM CaCl_2_, 10 mM Glucose) at 2-3 mL/min. For pulse labeling of ecDHFR in living cells, cell permeable TMP-hexachlorofluorescein (TMP-HEX, active motif) was applied at a concentration of 100 nM for 2 min followed by quick washout. To competitively block the labeling, DMSO (control) or 10 μM TMP was applied to the imaging solution (final concentration of DMSO was 0.1% [v/v]). Acquired image stacks were analyzed with NIH ImageJ/Fiji. After brief image registration, region of interests (ROI) were generated based on EGFP images by thresholding, and were used for quantification. After extraction of the average fluorescence intensity in each ROI, normalized fluorescence change (ΔF/F0) was plotted and used for quantitative analysis. In this experimental condition, TMP-HEX efficiently led to bright labeling of cells expressing the recombinant ecDHFR fused to EGFP but not control cells transfected with EGFP alone (**Figure S8A**). Fluorescence intensity reached a plateau within ~15 min, but it remained at a background level when TMP-HEX was mixed with an excess of unconjugated compound, supporting the saturability and specificity of TMP-HEX binding (**Figure S8B, 8C**).

### Two-photon microscope

*In vivo* labeling kinetics of ecDHFR reporter with TMP-HEX in living animal brain was analyzed by two-photon laser scanning microscopy (*44*). We used 900-nm wavelength pulse laser to elicit excitation of fluorophores. A single image plane was acquired at a fixed size (1,024 x 1,024 pixels square) with a depth of 0.2-0.4 mm from the surface. For simultaneous imaging of EGFP and TMP-HEX, emission signals were separated by beam splitter and detected with band-pass filters for green (525/50 nm) and red (610/75 nm), respectively. For kinetic analysis of DD-EGFP stabilization by TMP administration, TMP (Sigma, T7883) dissolved in 10% DMSO / normal saline was prepared and immediately used for i.p. injection at a final concentration of 100 mg/kg. Time course changes of ecDHFR-d2Venus fluorescence in response to air-puff stimulation of whiskers or chemogenetic activation of hM3-Dq with 10 mg/kg CNO i.p. injection were performed according to previously described methods (*14, 45*).

### Animal PET

PET scans were conducted with microPET Focus220 system (Siemens Medical Solutions USA, Malvern, USA) as described previously (*46*). Briefly, animals were anesthetized with 1.0-3.0% isoflurane during all PET procedures. For injection of PET tracers, dental needles conjugated to polyethylene tubing were inserted into the tail vein of mice and SURFLO F&F (26G, Terumo Corp., Tokyo, Japan) was inserted into the marmoset saphenous vein. After intravenous bolus injection of radioactive ligands (mice: 42.6±5.6 MBq for [^11^C]TMP, 40.7±13.2 MBq for [^18^F]FE-TMP, 37.0±1.3 MBq for [^11^C]GF120918, 32.3±6.1 MBq for [^11^C]verapamil, 25.5±15.8 MBq for [^18^F]MNI659, or 19.2±0.7 MBq for [^11^C]PBB3; marmoset: 112.7 MBq for [^18^F]FE-TMP), dynamic emission scanning with 3D acquisition mode was conducted for 60 min ([^11^C]PBB3), 90 min ([^11^C]TMP, [^18^F]FE-TMP [^11^C]GF120918, [^11^C]verapamil, and [^18^F]MNI659) or 180 min ([^18^F]FE-TMP), respectively. Data were histogrammed into a 3D set of sonograms, and images were reconstructed by either maximum a posteriori method or filtered back-projection using a Hanning filter cut-off at a Nyquist frequency (0.5 mm^-1^). All captured image data were subsequently analyzed using PMOD software (PMOD Technologies, Zurich, Switzerland). For spatial alignment of PET images, template MR images generated previously (*47*) were used in this study. For quantification of radioactive signal during PET scan, VOIs were manually placed on the image fields. In the analysis of P0-injected mouse brains, we placed VOIs in a region adjacent to the corpus callosum and hippocampus, which contain a maximum radioactive signal, and placed the reference in the striatal region with background signal. In the analysis of locally injected adult mouse brains, we manually set the same size of VOIs in the somatosensory-motor cortical region on both ipsilateral and contralateral sides. Average radioactive signal in each VOI was calculated as % standardized uptake value (%SUV), representing the % injected dose per cm^3^ volume compensated by body weight. PET analysis of monkey was performed as described previously (*15*).

At an initial stage of this study, we radiosynthesized [^11^C]TMP and characterized its performance in a PET analysis of mice that expressed ecDHFR-EGFP or red fluorescent protein, tdTomato, as a control generated by a single AAV injection into cerebral lateral ventricles at birth. At 2 months of age, mice were given [^11^C]TMP via tail vein bolus injection and subjected to dynamic PET scans. While animals carrying tdTomato exhibited minimal accumulation of [^11^C]TMP radioactivity, mice expressing ecDHFR-EGFP had strong radiosignals in the brain (**Figure S2A-S2A**). The PET imaging contrast for ecDHFR was profoundly improved when mice were given [^18^F]FE-TMP, and this was primarily attributed to a much faster washout of unbound [^18^F]FE-TMP from the brain after initial uptake (**Figure S2D-S2F**).

### Histology

Mice were deeply anesthetized and sacrificed by 4% PFA/PBS perfusion and fixation, and brains were subsequently dissected and further fixed in 4% PFA/PBS for 3 h at room temperature. For acquisition of whole brain imaging, fixed brains were imaged using M205FA fluorescence stereoscopic microscopy (Leica). After cryo-protection in PBS containing 20% sucrose, brains were embedded and frozen in OCT compound (SaKuRa), and 20-μm fixed frozen sections were prepared by cryostat. Alternatively, 100-μm free-floating coronal sections were sliced with a Leica VT1200S vibratome. All sections were blocked and permeabilized in PBS supplemented 4% BSA, 2% horse serum, and 0.25% Triton X-100 at room temperature, incubated with primary antibodies, and then probed with secondary antibodies labeled Alexa488, 546, or 633. After extensive wash-off of antibodies, nuclei were counter-stained with DAPI (Invitrogen) and mounted on glass slides. For postmortem verification of reporter distribution in the marmoset brain, the monkey was anesthetized and sacrificed by 4% PFA/PBS perfusion and fixation. Dissected brain tissue was cryo-protected in 30% sucrose/PBS followed by sectioning by cryostat. Slices of 20-μm thickness were then mounted on glass slides and fluorescence images were captured. For capturing low-magnification images of entire brain slices, sections were imaged using BZ-X710 fluorescence microscopy (Keyence) with x10 objective (NA 0.45). For high-magnification images to assess detailed cell type and/or structure, sections were analyzed by FV1000 laser scanning confocal microscopy (Olympus) with either x40 (NA 1.30) or x60 (NA 1.35) oil immersion objectives.

### *In vitro* binding assay

*In vitro* binding affinity of [^11^C]TMP ligand or [^18^F]FE-TMP ligand against ecDHFR reporter and MNI659 against PDE10 variants in HEK293T cell lysate was assessed as described previously (*48*). To assay radioligand binding with homologous blockade, these homogenates (50 μg protein) were incubated with 5 nM [^11^C]TMP, 1 nM [^18^F]FE-TMP or 1 nM [^18^F]MNI659 in the presence of unlabeled ligands at varying concentrations ranging from 10^-11^-10^-6^ M in Tris-HCl buffer, pH7.4, for 30 min at room temperature. These radioligand concentrations were determined in consideration of the radioactive half-lives of ^11^C (~20.4 min) and ^18^F (~109.8 min). Samples were run in duplicates or triplicates. Inhibition constant (Ki) and percentage of displacement were determined by using non-linear regression to fit a concentration-binding plot to one-site binding models derived from the Cheng-Prusoff equation with Prism software (GraphPad), followed by F-test for model selection. Dissociation constant (Kd) was calculated using this function: Kd = Ki = IC50 – [radioligand concentration].

For in vitro autoradiography, fixed frozen brain sections from mice expressing ecDHFR-EGFP or tdTomato as control were pre-incubated in 50 mM Tris-HCl buffer, pH7.4, containing 5% ethanol at room temperature for 30 min, and incubated in 50 mM Tris-HCl buffer, pH7.4, containing 5% ethanol and [^18^F]FE-TMP (10 mCi/L) at room temperature for 30 min. Sections were then rinsed with ice-cold Tris-HCl buffer containing 5% ethanol twice for 2 min, and dipped into ice-cold water for 10 sec. Sections labeled with [^18^F]FE-TMP were subsequently dried with warm air and exposed to an imaging plate (BAS-MS2025; Fuji Film). The imaging plate was scanned with a BAS-5000 system (Fuji Film) to acquire autoradiograms. To obtain evidence supporting our *in vivo* imaging data on the specificity of fluorescent and radioactive TMP analogs for ecDHFR, *in vitro* heterologous blockade of reporter labeling by a well characterized ecDHFR ligand, TMP, was analyzed. Readouts of both fluorescence microscopy and autoradiography consistently demonstrated that, in brain sections derived from mice carrying ecDHFR-EGFP, TMP-HEX (**Figure S9A**) and [^18^F]FE-TMP (**Figure S9B**) were selectively retained in areas of transgene expression, and their detection was strongly suppressed in the presence of saturating amounts (100 μM) of TMP.

### Metabolite analysis

To analyze the radioactive metabolite of [^18^F]FE-TMP in plasma and brain of mice, blood was collected from the heart at 5, 15, 30, 60 and 90 min after intravenous injection of [^18^F]FE-TMP (100–250 MBq) under anesthesia, followed by quick removal of the brain. Blood samples were centrifuged at 15,000 rpm for 3 min at 4°C. Plasma (200 μL) was transferred to a test tube containing acetonitrile (200 μL). The mixture was vortexed and centrifuged at 15,000 rpm for 3 min at 4°C for deproteinization. The brain was homogenized in ice-cold saline. The homogenate (500 μL) was mixed with acetonitrile (500 μL) and centrifuged at 15,000 rpm for 3 min at 4°C for deproteinization. More than 93% of radioactivity in the tissues was extracted in the supernatants. Radioactive components in the supernatants were analyzed using an HPLC system equipped with a high-sensitive positron detector (Ohyo Koken Kogyo Co. Ltd., Tokyo, Japan). The HPLC system and conditions were as follows: pump, PU-2089 plus (Jasco); UV detector, UV-2075 (Jasco); the high-sensitive positron detector; column, Capcell Pak C18 AQ (Shiseido Co., Ltd, Tokyo, Japan), 5 μm, 10 i.d. × 250 mm; mobile phase, MeCN/0.1% phosphoric acid = 1/9; flow rate 4 mL/min; retention time, 12 min.

Two major radiometabolites (M1 and M2) were observed (**Figure S10A**). M2 was present in plasma; its fraction reached a plateau at 15 min after i. v. injection of [^18^F]FE-TMP and did not undergo efficient transfer to the brain (**Figure S10C**). In contrast, M1 was a major radioactive metabolite in the brain, and its fraction gradually increased over 90 min after i. v. injection of [^18^F]FE-TMP (**Figure S10B**). Since TMP undergoes *O*-demethylation and the preliminary metabolite analysis described above indicated that metabolite M1 was highly hydrophilic, M1 was expected to be [^18^F]fluoroacetic acid ([^18^F]FAcOH; see **Figure S10D**) (*21, 39*). Then, we compared the retention time of M1 with that of [^18^F]fluoroacetic acid, which was generated by the incubation of ethyl [^18^F]fluoroacetate in the plasma of mouse at 37°C for 30 min (Mori et al., 2009), using a normal phase HPLC (column, Cosmosil HILIC (Nacalai Tesque Inc., Kyoto, Japan), 5 μm, 4.6 i.d. × 150 mm; mobile phase, MeCN/AcONH4 (10 mM, pH 7.0) = 1/1; flow rate, 1 mL/min) and a reverse phase HPLC (column, Capcell Pak C18 AQ (Shiseido Co., Ltd, Tokyo, Japan), 5 μm, 10 i.d. × 250 mm; mobile phase, MeCN/0.1% phosphoric acid (0–5 min, 9/1; 5–10 min, 9/1 to 1/1; 10–13 min, 1/1; 13–15 min, 1/1 to 9/1); flow rate, 4 mL/min). As expected, the retention time of M1 was identical to that of [^18^F]fluoroacetic acid in these analysis conditions. Estimated accumulation of [^18^F]FAcOH in the brain was relatively slow (**Figure S10E**), and this metabolite should not react with ecDHFR in consideration of its chemical structure.

### Statistics

Statistical significance of the data was analyzed with GraphPad Prism software (Graphpad Inc.). Unpaired Student’s t-test or Mann-Whitney test was used for comparison of two-group data. For comparison of multiple groups, data were analyzed by one-way ANOVA post hoc Dunnett test or Tukey-Kramer test. For analysis of time series data, statistical significance was determined by two-way, repeated measure ANOVA followed by Bonferroni’s multiple comparisons test.

## Data availability

The data that support the findings of this study are available from the corresponding author upon reasonable request.

## Acknowledgements

We thank Dr. Atsushi Miyawaki (RIKEN, Saitama, Japan) and Dr. Haruo Kasai (Tokyo University, Tokyo, Japan) for discussions and comments on the manuscript. We thank Drs. Tetsuo Yamamori and Masanari Ohtsuka (RIKEN, Saitama, Japan) for sharing materials. We thank Shoko Uchida, Kaori Yuki, and Katsushi Kumada for technical assistance. This study was supported in part by JST CREST Grant Number JPMJCR1652 (M.H.), AMED Grant Number JP19dm0207072 (M.H.) and JP19dm0107146 (T.M.), and JSPS KAKENHI Grant Number 18H04752 (M.S.) and 18K07777 (M.S.).

## Author contributions

M.S. and M.H. conceived the study; M.S. generated expression vectors and recombinant viruses, and performed biochemical and histological analyses; M.S., H.T., M.T., Y.S., Y.T., and N.S. carried out *in vitro* and *in vivo* fluorescence imaging studies; M.O. performed autoradiography and *in vitro* binding assays; J.M, T.M., M.O., and T.K. analyzed TMP metabolites; M.-R.Z. and M.F. supervised chemical design, synthesis, and radioactive probing; M.S., M.T., C.S. and Y.T. performed PET and data analysis; K.M., Y.N., and T.M. performed PET and histological analysis of the primate. M.H, A.M, N.S, T.S. and M.S. wrote the manuscript.

