## Supplemental Materials for "Genetically targeted reporter imaging of deep neuronal network in the mammalian brain"

**Title**

Figure S1

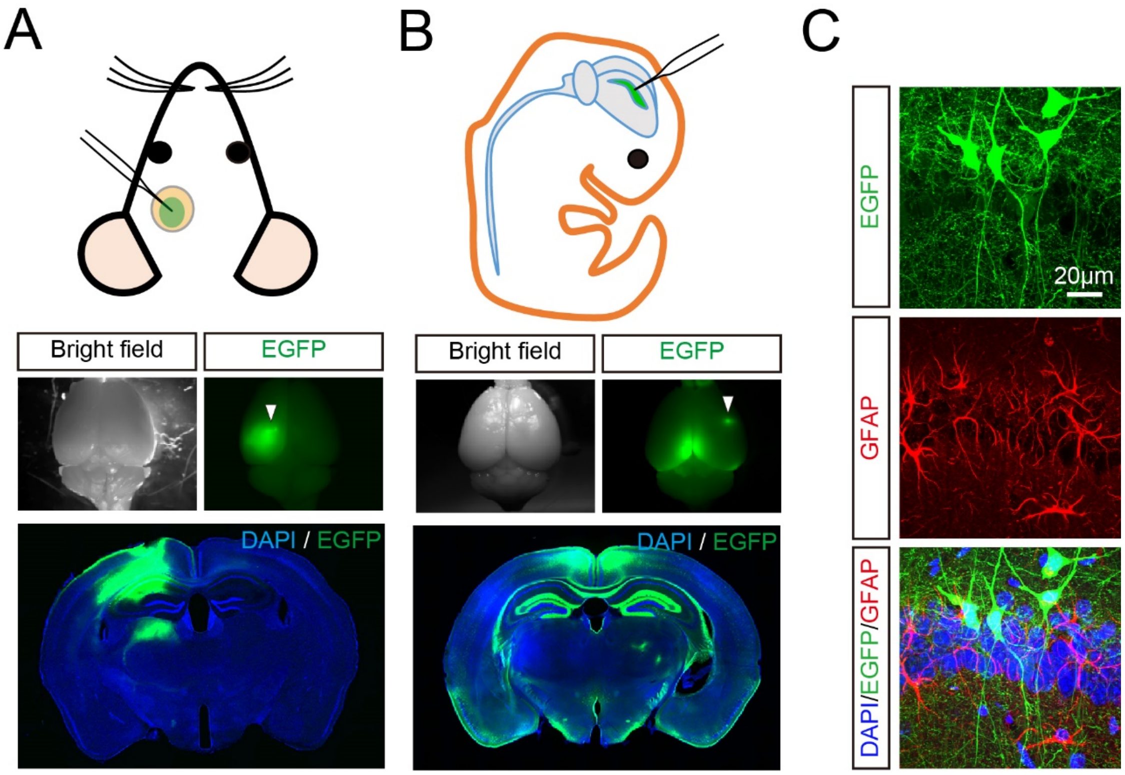

A
B
C
Bright field
EGFP
Bright field
EGFP
DAPI / EGFP
DAPI / EGFP
EGFP
GFAP
DAPI/EGFP/GFAP
20μm

**Figure S1. Comparative assessment of transgene expression by AAV injection into rodent brain with two different protocols**

**A**) Schematic diagram of AAV injection into one side of the somatosensory cortex with cranial window in adult mouse brain. EGFP expression was introduced under control of synapsin promoter. Spatial distribution of EGFP was analyzed in the whole brain (upper) and coronal brain slices (lower). Note that EGFP expression was highly restricted to the injection site in this protocol.

**B)** Schematic diagram of AAV injection into one side of lateral cerebral ventricle at postnatal day 0. Spatial distribution of EGFP was analyzed in the whole brain (upper) and coronal slices (lower). Note that high-level expression of EGFP is observed in circumferentially arranged area of the ventricle system including the corpus callosum, hippocampus, and parietal cortex.

**C)** Neuron-specific expression of EGFP under control of the synapsin promoter is verified by immunostaining using anti-GFAP antibody. Representative images of neurons (green) and astrocytes (red) with DAPI staining (blue) in hippocampal CA1 captured by confocal microscopy are displayed.

#### Figure S2

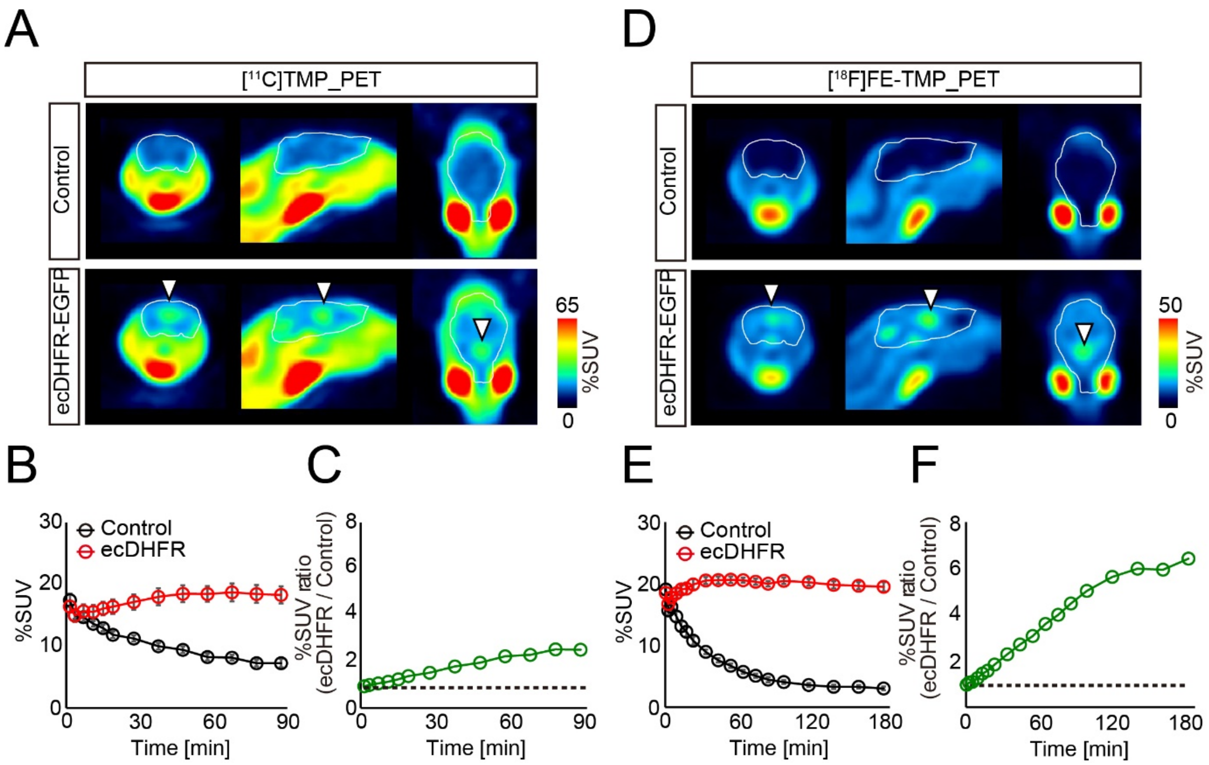
A
[11C]TMP_PET
Control
ecDHFR-EGFP
65
%SUV
0
D
[18F]FE-TMP_PET
Control
ecDHFR-EGFP
50
%SUV
0
B
Control
ecDHFR
%SUV
Time [min]
C
%SUV ratio (ecDHFR / Control)
Time [min]
E
Control
ecDHFR
%SUV
Time [min]
F
%SUV ratio (ecDHFR / Control)
Time [min]

**Figure S2. PET imaging of ecDHFR in mouse brain with radioactive TMP analogs**

Mice expressing ecDHFR-EGFP or tdTomato (as control) from AAVs in the forebrain were subjected to PET scans after peripheral administration of the radioactive TMP analogs $[^{11}C]$TMP and $[^{18}F]$FE-TMP (40 MBq/mouse).

**A)** Representative PET images (coronal, sagittal and horizontal sections from left) generated by averaging dynamic scan data at 0-90 min after i.v. injection of $[^{11}C]$TMP. Arrowheads indicate areas of accumulation of radioactive ligand in animals carrying ecDHFR-EGFP (lower). White lines mark whole brain area as determined by MRI.

**B)** $[^{11}C]$TMP labeling kinetics. Volumes of interest (VOI) of fixed sizes were manually placed on the paraventricular region exhibiting high-level radioactive signals. Data from control ($n = 7$) and ecDHFR-EGFP-expressing mice ($n = 7$) were plotted as Mean ± SEM. $F(1, 12) = 22.05$; $p < 0.01$ (two-way ANOVA).

**C)** Ratios of averaged $[^{11}C]$TMP radioactive signals in ecDHFR versus control brains.

**D)** Representative PET images (coronal, sagittal and horizontal sections from left) generated by averaging dynamic scan data at 0-180 min after i.v. injection of $[^{18}F]$FE-TMP. Arrowheads indicate areas of accumulation of radioactive ligand in animals carrying ecDHFR-EGFP (lower). White lines mark whole brain.

**E)** $[^{18}F]$FE-TMP labeling kinetics. VOI analysis was performed as described in panel **c**. Data from control ($n = 6$) and ecDHFR-EGFP-expressing mice ($n = 6$) were plotted as Mean ± SEM. $F(1, 10) = 326.1$; $p < 0.01$ (two-way ANOVA).

**F)** Ratios of averaged $[^{18}F]$FE-TMP radioactive signals in ecDHFR versus control brains.

### Figure S3

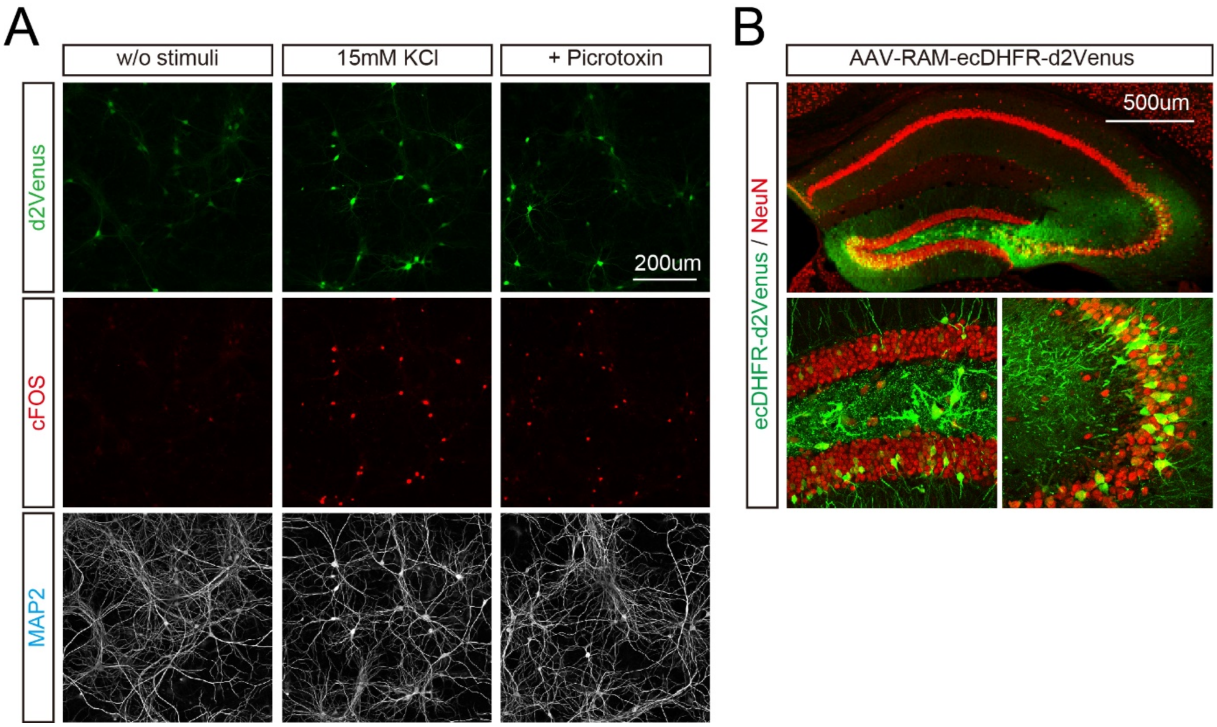


**Figure S3. Efficient labeling of activated neurons with RAM promoter *in vitro* and *in vivo***

**A)** DIV21 neuronal culture expressing d2Venus under control of RAM promoter was incubated with 15 mM KCl (middle) or picrotoxin (right) for 6 h. Fixed neurons were immunostained with anti-cFOS and anti-MAP2 antibodies and analyzed by confocal fluorescence microscopy. Representative images demonstrate enhanced signals of d2Venus fluorescence and cFos immunoreactivity in neurons after chemical stimuli.

**B)** AAV-RAM-ecDHFR-d2Venus and AAV-CAG-M3-DREADD were co-injected into one side of the hippocampus. The animal was sacrificed one week after 0.3 mg/kg CNO i.p. injection for PET analysis, and distribution of ecDHFR-d2Venus in brain slices was assessed by confocal microscopy.

### Figure S4

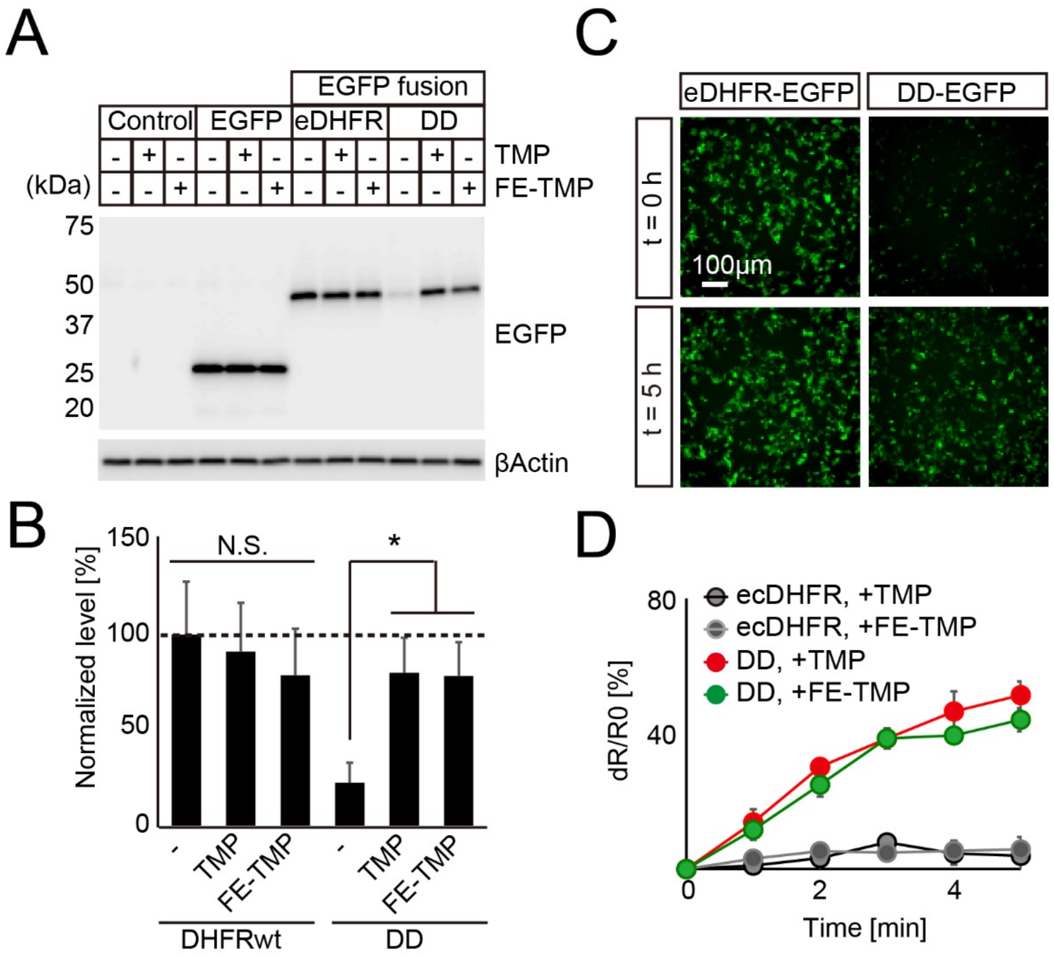

A
EGFP fusion
Control
EGFP
eDHFR
DD
TMP
FE-TMP
(kDa)
75
50
37
25
20
EGFP
βActin
C
eDHFR-EGFP
DD-EGFP
t = 0 h
t = 5 h
100μm
B
Normalized level [%]
150
100
50
0
N.S.
*
-
TMP
FE-TMP
DHFRwt
DD
D
ecDHFR, +TMP
ecDHFR, +FE-TMP
DD, +TMP
DD, +FE-TMP
dR/R0 [%]
80
40
0
2
4
Time [min]

**Figure S4. Stabilization of destabilized ecDHFR mutant by TMP analogs**

**A-D)** Cultured HEK293 cells expressing ecDHFR-EGFP or DD-EGFP were incubated with 10 μM TMP or FE-TMP and analyzed by immunoblotting and fluorescence microscopy.

**A)** Protein stabilization of wild-type ecDHFR and DD fused to EGFP were analyzed by immunoblotting with anti-GFP chicken IgY. For loading control, protein levels of β-actin are assessed with specific antibodies. Note that incubation with TMP and FE-TMP did not affect protein levels of wild-type ecDHFR-EGFP but remarkably induced stabilization of DD during the observation for 24 h.

**B)** Quantitative analysis of immunoblot data. Data from three independent experiments are shown as Mean ± SD. $F(5, 12) = 4.746$; $*p < 0.05$ (one-way ANOVA followed by Dunnett post-hoc test).

**C)** Time course analysis of DD-EGFP stabilization by TMP in HEK293T cells. Plasmid DNA vectors encoding either wild-type ecDHFR-EGFP or DD-EGFP were transfected into cells with control vectors coding mCherry. From 30 h after transfection, protein stabilization of DD-EGFP by 10 μM ligand addition to the culture medium was monitored hourly. Note that only DD-EGFP fluorescence signal is specifically upregulated by TMP addition.

**D)** Ratiometric quantitative analysis of fluorescence signals of EGFP relative to mCherry fluorescence. Changes of EGFP-to-mCherry ratio at each time point is shown as % of the ratio at $t = 0$. Data from three independent experiments are displayed as Mean ± SD. Note that TMP and FE-TMP possess similar ability to stabilize DD proteins in this condition.

### Figure S5

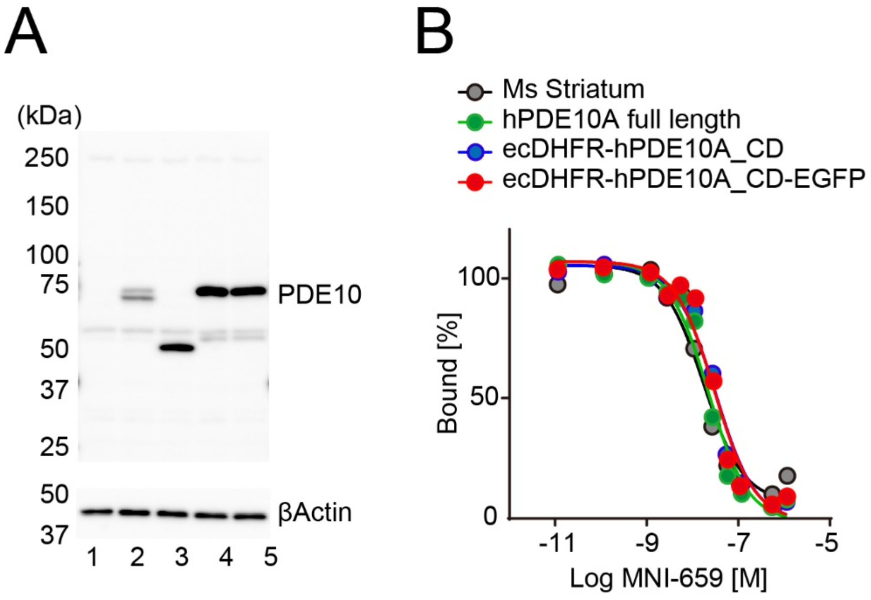


**Figure S5. Analyses of ecDHFR reporters fused with cyclic phosphodiesterase (PDE10A)**

**A)** Expression of indicated constructs in cultures of HEK293 cells. Protein extracts were analyzed by immunoblotting with antibodies against PDE10 or β-Actin (as loading control). Lane 1: vector control, 2: full-length PDE10A, 3: ecDHFR-PDE10A_CD, 4: ecDHFR-PDE10A_CD-EGFP, 5: ecDHFR-PDE10A(D674A)_CD-EGFP.

**B)** Binding of [$^{18}$F]MNI659 to various recombinant PDE10A reporters was assessed by competition assay with various concentrations of non-labeled MNI659 compound. Mouse striatum homogenate, which contains endogenous PDE10A, was used as positive control. Apparent Bmax (Maximum Binding) was set at 100%.

### Figure S6

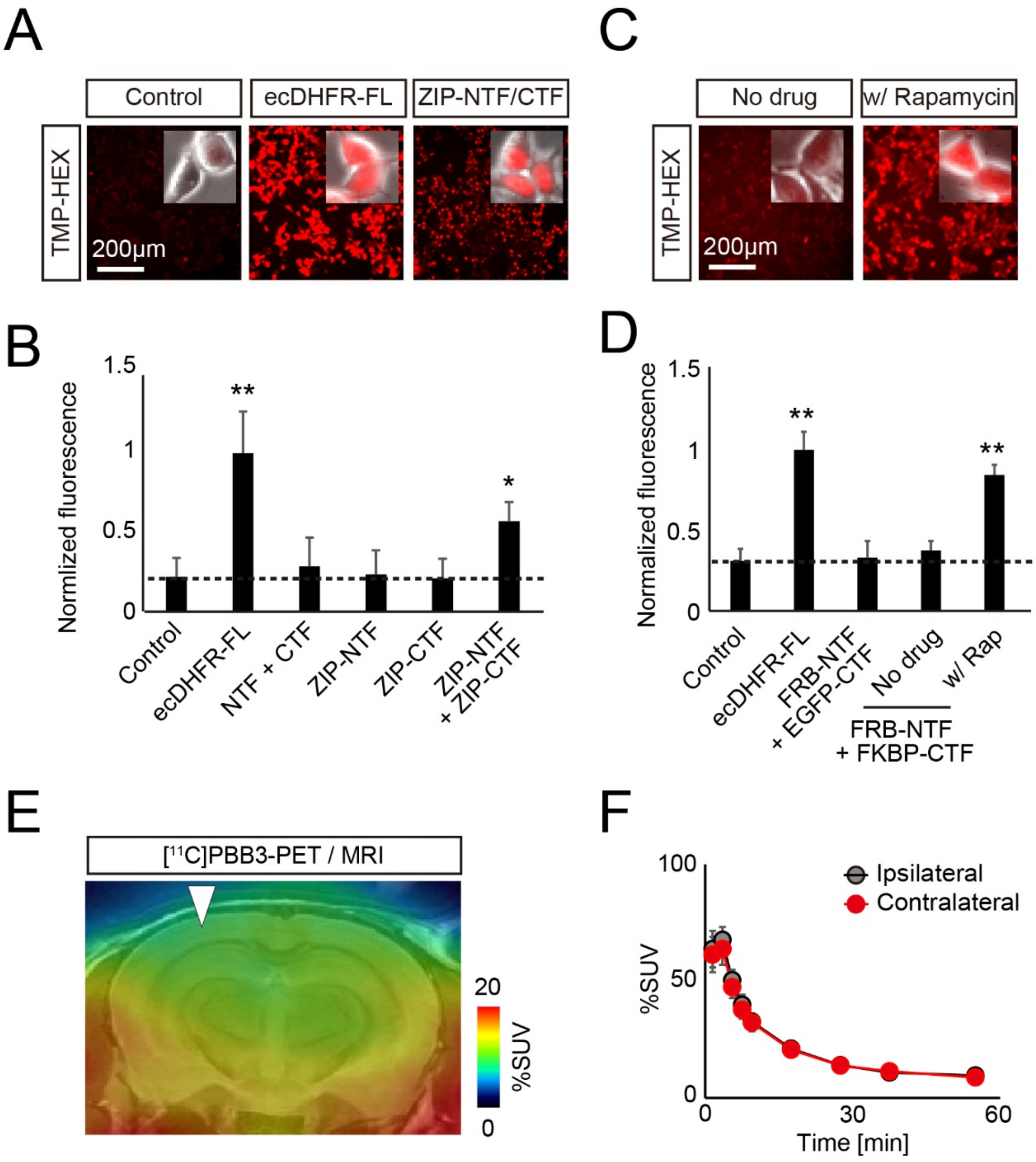
A
Control
ecDHFR-FL
ZIP-NTF/CTF
TMP-HEX
200μm
B
Normlized fluorescence
1.5
1
0.5
0
**
*
Control
ecDHFR-FL
NTF + CTF
ZIP-NTF
ZIP-CTF
ZIP-NTF + ZIP-CTF
C
No drug
w/ Rapamycin
TMP-HEX
200μm
D
Normalized fluorescence
1.5
1
0.5
0
**
**
Control
ecDHFR-FL
FRB-NTF + EGFP-CTF
No drug
w/ Rap
FRB-NTF + FKBP-CTF
E
[11C]PBB3-PET / MRI
20
0
%SUV
F
100
50
0
%SUV
Ipsilateral
Contralateral
0
30
60
Time [min]

**Figure S6. Design and characterization of protein fragment complementation assay using a split DHFR system**

**A-D)** Cultured HEK293 cells expressing various constructs were incubated with 100 nM TMP-HEX and analyzed by fluorescent microscopy. TMP-HEX efficiently labeled cultured cells co-expressing NTF and CTF conjugated to a self-assembling leucine zipper (ZIP) motif, or a rapamycin-dependent heterodimerization motifs FKBP12-rapamycin binding domain (FRB) and FK506 binding protein (FKBP) in the rapamycin-dependent manner, indicating that the cDHFR-PCA system functions under these conditions as designed.

**A)** Representative images illustrate a selective retention of TMP-HEX in cells carrying the full-length ecDHFR (as positive control) or a combination of ZIP-tagged ecDHFR NTF and CTF. Insets demonstrate high magnification fluorescence images overlaid with phase contrast pictures of individual cells. Note that ZIP-tagged ecDHFR NTF and CTF preferentially localize in nucleus.

**B)** Normalized fluorescence intensities of TMP-HEX in cells transfected with indicated constructs. Mean value of full-length ecDHFR (ecDHFR-FL) was set as 1. Note that only ecDHFR-FL and the split protein that reassembles via ZIP interaction retain the marker above background levels. Data from six independent experiments are plotted as Mean ± SD. $F(5, 28) = 21.12$; $^{**}p < 0.01$ (one-way ANOVA followed by Dunnett post-hoc test).

**C)** HEK293 cells expressing various constructs were incubated with 100 nM TMP-HEX with or without 500 nM rapamycin (LC Laboratories) and imaged by fluorescence microscopy. Representative images illustrate selective labeling of TMP-HEX in cells co-expressing ecDHFR NTF and CTF tagged with FRB and FKBP in the presence of rapamycin. Insets demonstrate high magnification fluorescence photomicrographs overlaid with phase contrast images.

**D)** Rapamycin (Rap) induced FRB-FKBP interaction was assessed as TMP-HEX labeling efficiency in cells expressing various combinations of ecDHFR-NTF and CTF fragments. Fluorescence intensities were normalized by mean value of labeling of full-length ecDHFR (ecDHFR-FL) with TMP-HEX. Data from four independent experiments are presented as

Mean ± SD. $F(4, 15) = 55.08$; $^{**}p < 0.01$ (one-way ANOVA followed by Dunnett post-hoc test).

**E)** A representative coronal PET image captured with [$^{11}$C]PBB3, a potent PET tracer for aggregated tau fibrils, in mice expressing TRD-NTF and TRD-CTF by AAV injection into one side of the somatosensory cortex. Averaged image of dynamic scan data at 0-60 min after i. v. injection of [$^{11}$C]PBB3 is shown. A template MRI image was overlaid for spatial alignment of the PET image. Arrowhead indicates injection site.

**F)** Kinetics of [$^{11}$C]PBB3 in mouse brain during 60-min dynamic PET scan. VOIs were manually placed on ipsilateral and contralateral cortical areas for quantification. Data from four mice expressing TRD-NTF and TRD-CTF are plotted as Mean ± SEM. Note there is no significant difference in radioactive signals of [$^{11}$C]PBB3 between ipsilateral and contralateral sides.

### Figure S7

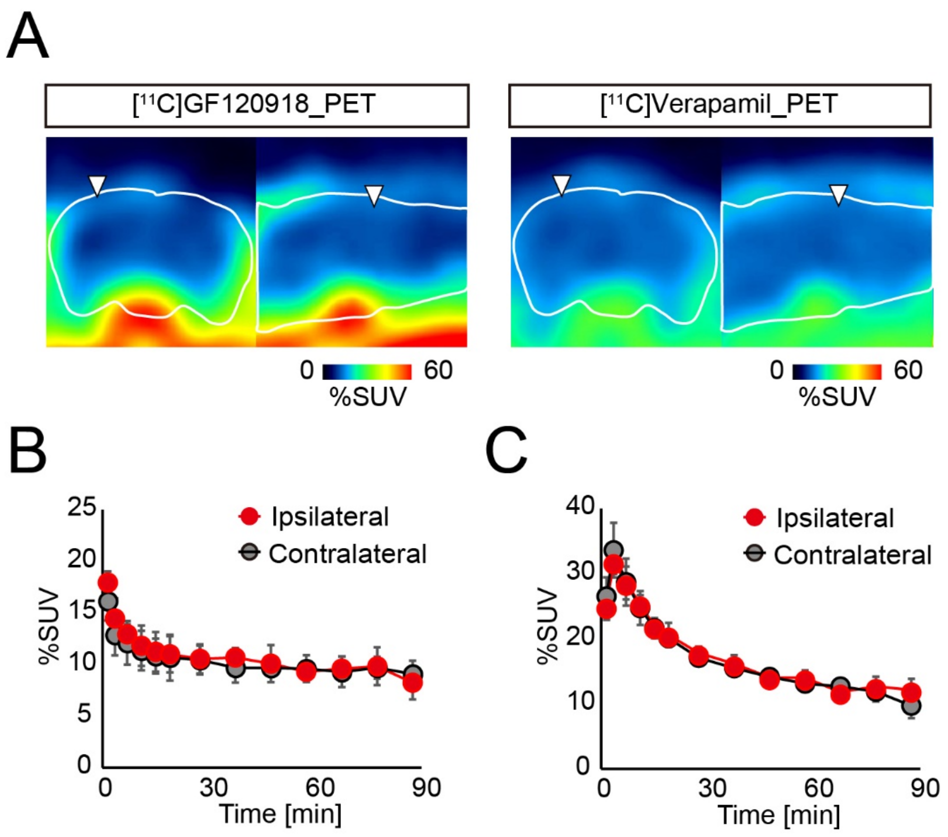
A
[11C]GF120918_PET
0
60
%SUV
[11C]Verapamil_PET
0
60
%SUV
B
Ipsilateral
Contralateral
%SUV
25
20
15
10
5
0
0
30
60
90
Time [min]
C
Ipsilateral
Contralateral
%SUV
40
30
20
10
0
0
30
60
90
Time [min]

**Figure S7. BBB integrity in mouse brains after local AAV delivery**

BBB integrity in brains of AAV-injected mice was assessed by PET scans with the radiolabeled substrates for p-glycoprotein [$^{11}$C]GF120918 and [$^{11}$C]verapamil (1 month post-injection).

**A)** Representative PET images (coronal and sagittal sections from left) generated by averaging dynamic scan data at 0 – 90 min after i.v. injection of [$^{11}$C]GF120918 (left) and [$^{11}$C]verapamil (right). White lines mark whole brain area as determined by MRI. Arrowheads indicate areas of AAV delivery. Scale bar represents %SUV.

**B)** Time-radioactivity curves in ipsilateral (red symbols) and contralateral (black symbols) cortices after administration of [$^{11}$C]GF120918. Note that AAV inoculation did not induce alteration of radiotracer retention in ipsilateral versus contralateral cortex. Data are Mean ± SD (n = 5 mice).

**C)** Time-radioactivity curves in ipsilateral (red symbols) and contralateral (black symbols) cortices after administration of [$^{11}$C]verapamil. Data are Mean ± SD (n = 4 mice).

### Figure S8

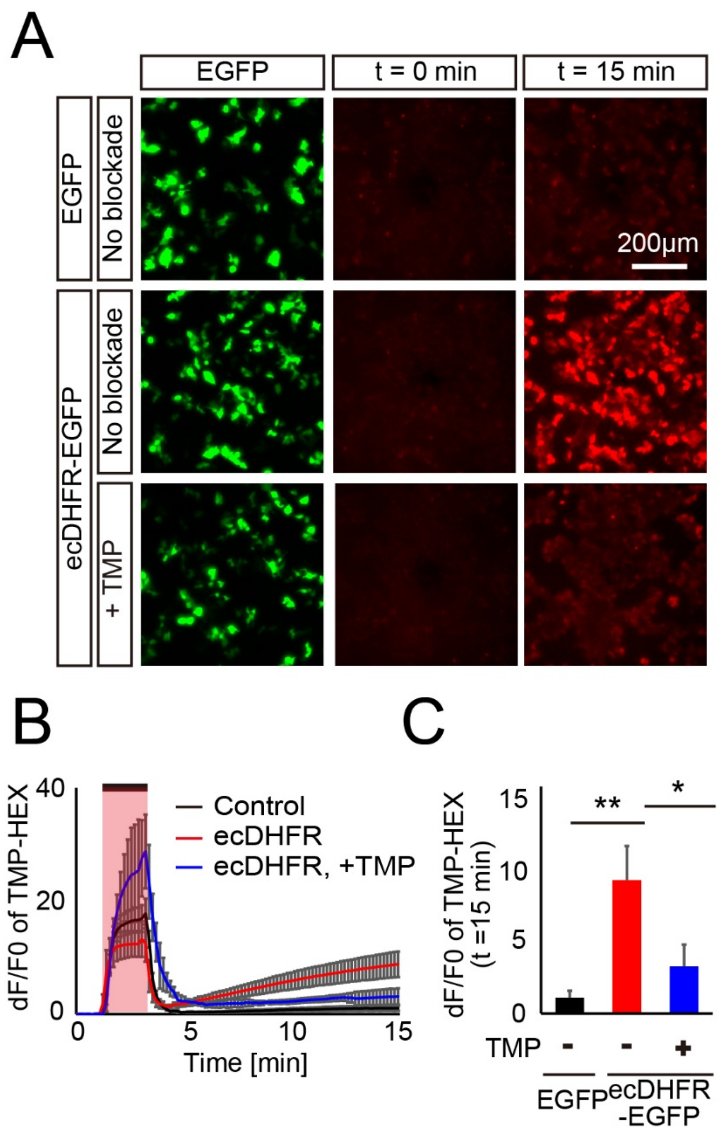
A
EGFP
t = 0 min
t = 15 min
EGFP
No blockade
200μm
ecDHFR-EGFP
No blockade
+ TMP
B
dF/F0 of TMP-HEX
40
20
0
Control
ecDHFR
ecDHFR, +TMP
0
5
10
15
Time [min]
C
dF/F0 of TMP-HEX
(t =15 min)
15
10
5
0
**
*
TMP
-
-
+
EGFP
ecDHFR
-EGFP

**Figure S8.** ***In vitro*** **fluorescent labeling of ecDHFR with TMP-HEX**

Cultured HEK293T cells expressing EGFP or ecDHFR-EGFP were imaged in time-lapse mode before, during, and after transient perfusion of 200 nM TMP-Hexachlorofluorescein (TMP-HEX). The ligand was applied for 2 min followed by a wash-out.

**A)** Representative images show selective retention of TMP-HEX (red) in cells carrying ecDHFR-EGFP (middle). Note that labeling is strongly suppressed in the presence of excess conventional TMP (10 μM, lower).

**B)** TMP-HEX fluorescence intensities during time-lapse image sessions, plotted as dF/F0 ratios. The bar marks the 2-min window of TMP-HEX perfusion. Data from three independent experiments are indicated as Mean ± SD. At 5-15 min, $F(2,6) = 23.08$; $p < 0.01$ (two-way, repeated measure ANOVA).

**C)** dF/F0 ratios of TMP-HEX fluorescence intensities at the endpoint of the assay (t = 15 min). Data from three independent experiments are indicated as Mean ± SD. $F(2,6) = 19.80$; *$p < 0.05$, **$p < 0.01$ (one-way ANOVA followed by Tukey-Kramer post-hoc test).

### Figure S9

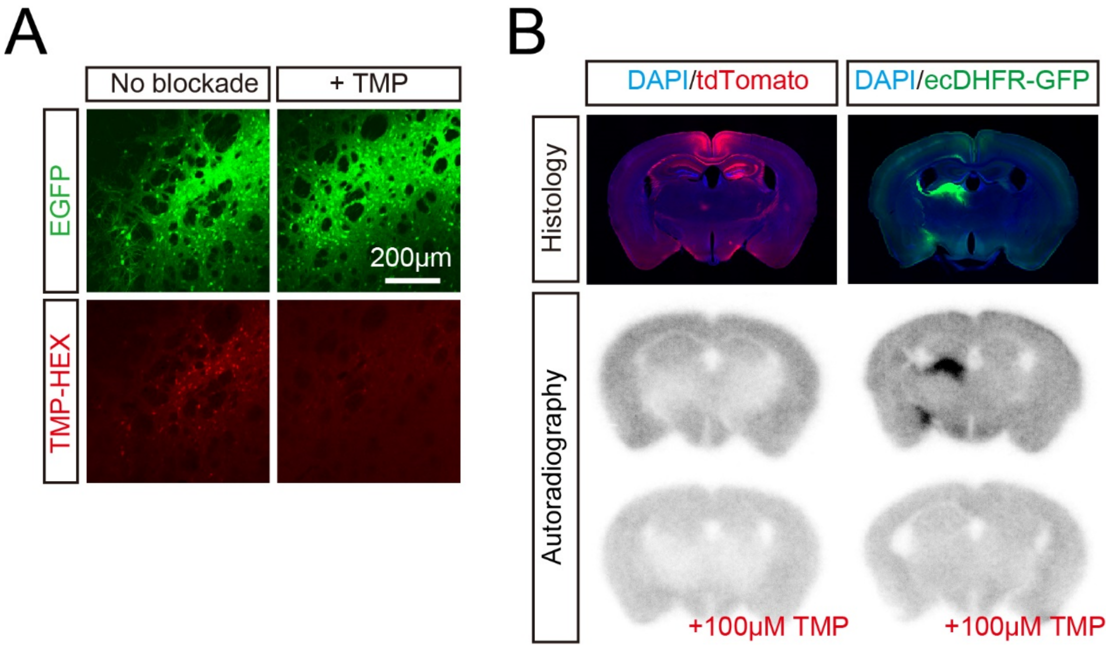


**Figure S9. *In vitro* validation of TMP-HEX and [$^{18}$F]FE-TMP for ecDHFR**

**A)** Representative images of striatal neurons expressing high-level ecDHFR-EGFP in fixed brain slices labeled with TMP-HEX. Note that incubation with excess amount of non-labeled TMP markedly inhibits fluorescence labeling.

**B)** *In vitro* autoradiography of mouse brain sections with [$^{18}$F]FE-TMP. Expression of transgenes in samples collected from mice treated with control and ecDHFR-EGFP vectors is fluorescently visualized with DAPI counterstaining (upper). Representative images of *in vitro* autoradiography using [$^{18}$F]FE-TMP demonstrate that this radioligand specifically labels brain areas overexpressing ecDHFR-EGFP but not tdTomato (middle), and that this radioligand binding to putative ecDHFR is profoundly blocked by an excess amount of non-labeled TMP (lower).

### Figure S10

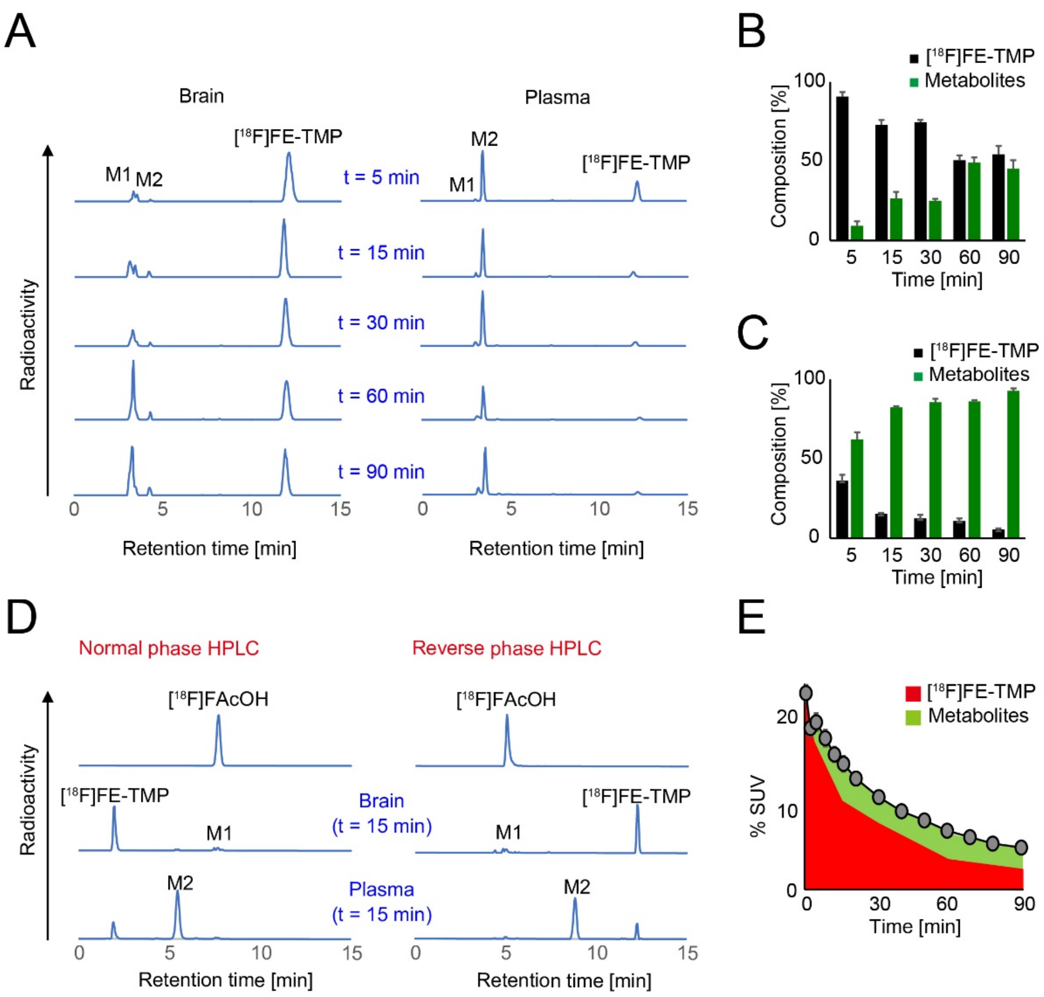

A
Brain
Plasma
[18F]FE-TMP
M1
M2
t = 5 min
t = 15 min
t = 30 min
t = 60 min
t = 90 min
Radioactivity
Retention time [min]
0 5 10 15
B
C
[18F]FE-TMP
Metabolites
Composition [%]
100 50 0
5 15 30 60 90
Time [min]
D
Normal phase HPLC
Reverse phase HPLC
[18F]FAcOH
Brain (t = 15 min)
Plasma (t = 15 min)
E
% SUV
20 10 0
0 30 60 90

**Figure S10. Characterization of [$^{18}$F]FE-TMP radioactive metabolites**

**A)** Representative reverse phase radio-HPLC charts showing [$^{18}$F]FE-TMP and its radiometabolites, M1 and M2, in brain (left) and plasma (right) of mice at 5, 15, 30, 60 and 90 min after i. v. injection of [$^{18}$F]FE-TMP. M1 and M2 are detected as major radiometabolites in brain and plasma, respectively.

**B-C)** Time course changes in the composition of [$^{18}$F]FE-TMP and its radiometabolites in brain (**B**) and plasma (**C**). Relative amounts of radiomaterials are indicated as % of total radioactivities at each time point.

**D)** Identification of a major metabolite, M1, by normal (left) and reverse (right) phase radio-HPLC. Retention times in HPLC charts were compared between radiosynthesized [$^{18}$F]fluoroacetate ([$^{18}$F]FAcOH) and radioactive metabolites derived from [$^{18}$F]FE-TMP in mouse brain and plasma at 15 min after i. v. administration of [$^{18}$F]FE-TMP. Retention time of [$^{18}$F]FAcOH was identical to that of M1, a major radiometabolite of [$^{18}$F]FE-TMP in brain.

**E)** Radioactivity derived from [$^{18}$F]FE-TMP (red area) and its radiometabolites (green area) in control cortex was calculated by applying temporal changes of their relative abundance (**B**) to the time-radioactivity curve shown in **Figure S2E**.
